# A megaplasmid family responsible for dissemination of multidrug resistance in *Pseudomonas*

**DOI:** 10.1101/630780

**Authors:** Adrian Cazares, Matthew P. Moore, Macauley Grimes, Jean-Guillaume Emond-Rhéault, Laura L. Wright, Pisut Pongchaikul, Pitak Santanirand, Roger C. Levesque, Joanne L. Fothergill, Craig Winstanley

## Abstract

Multidrug resistance (MDR) represents a global threat to health. Although plasmids can play an important role in the dissemination of MDR, they have not been commonly linked to the emergence of antimicrobial resistance in the pathogen *Pseudomonas aeruginosa*. We used whole genome sequencing to characterize a collection of *P. aeruginosa* clinical isolates from a hospital in Thailand. Using long-read sequence data we obtained complete sequences of two closely related megaplasmids (>420 kb) carrying large arrays of antibiotic resistance genes located in discrete, complex and dynamic resistance regions, and revealing evidence of extensive duplication and recombination events. A comprehensive pangenomic and phylogenomic analysis indicated that 1) these large plasmids comprise a family present in different members of the *Pseudomonas* genus and associated with multiple sources (geographical, clinical or environmental); 2) the megaplasmids encode diverse niche-adaptive accessory traits, including multidrug resistance; 3) the pangenome of the megaplasmid family is highly flexible and diverse, comprising a substantial core genome (average of 48% of plasmid genes), but with individual members carrying large numbers of unique genes. The history of the megaplasmid family, inferred from our analysis of the available database, suggests that members carrying multiple resistance genes date back to at least the 1970s.

**Funding:** This work was supported by the International Pseudomonas Genomics Consortium, funded by Cystic Fibrosis Canada [RCL]; and the Secretaría de Educación, Ciencia, Tecnología e Innovación (SECTEI), Mexico [AC].

## Introduction

The spread of antimicrobial resistance (AMR) is recognised as a key global challenge to human health (1). The so-called ESKAPE pathogens (*Enterococcus faecium, Staphylococcus aureus, Klebsiella pneumoniae, Acinetobacter baumannii, Pseudomonas aeruginosa* and *Enterobacter* sp) are the leading cause of nosocomial infections worldwide (2). In particular, *P. aeruginosa* causes a wide range of opportunistic infections (3), often associated with multidrug resistance (MDR) (4). Indeed *P. aeruginosa* (especially carbapenem resistant strains) has been highlighted by the World Health Organisation as one of the critical (Priority 1) pathogens associated with AMR (5).

Alongside the development of new drugs, it is crucial that we improve our understanding of the factors driving the evolution and spread of AMR. As with other processes of bacterial adaptation, the rapid spread of resistance is often facilitated by mobile genetic elements (MGE): entities that are adapted to moving DNA between replicons and between cells (6), including insertion sequences, transposons, integrons and conjugative plasmids, which can facilitate the spread of AMR genes to different host backgrounds (7). In contrast to pathogens from the enterobacteriaceae, the role of plasmids in the spread of MDR in *P. aeruginosa* is not well understood.

It is recognised that AMR can develop and spread rapidly in regions where antibiotics are inappropriately sold/used and freely available, and that hospital settings provide excellent opportunities for the dissemination of resistance. In Thailand, carbapenem-resistant *P. aeruginosa* is an increasing problem, especially in the context of hospital-acquired infections (8, 9). Previous studies have sought to characterise the genetic basis underlying carbapenem resistance amongst *P. aeruginosa* clinical isolates in Thailand (10), and globally (11), but the role of plasmids is poorly understood.

Whole genome sequencing can be a powerful epidemiological tool in diagnostic and public health microbiology (12, 13), but it has limitations that have restricted our understanding of the role of MGE. The use of short-read data from the most popular high-throughput platforms makes the assembly of plasmids, which often contain repetitive sequences, difficult (14). Long read sequencing, such as PacBio or Oxford Nanopore, can resolve the status of plasmids, allowing comparative analysis that can elucidate the routes by which AMR genes spread between lineages (15).

Here we describe a combination of long-read and short-read genome sequencing that allowed us to characterise a family of megaplasmids contributing to the spread of MDR *P. aeruginosa* in a hospital in Thailand. By surveying sequence databases, members of this megaplasmid family were also identified in isolates from multiple sources from around the world, including the environment, and in non-*aeruginosa* species of *Pseudomonas*. Related plasmids have a shared core genome, but vary in the carriage of AMR genes, identifying the megaplasmid as a vehicle for the consolidation and dissemination of resistance. Our study provides valuable insights into how MDR plasmids emerge from environmental reservoirs into a clinical setting.

## Materials and Methods

### Bacterial isolates

The 48 clinical isolates used in this study were acquired in 2013 from patients in Ramathibodi Hospital, Mahidol University, Bangkok, Thailand, and were associated with different types of infection (Supplementary Table S1).

### Antimicrobial susceptibility testing

Antimicrobial susceptibility testing was carried out according to the The European Committee on Antimicrobial Susceptibility Testing (EUCAST) guidelines (16). Briefly, isolates to be tested were cultured onto Columbia plates (overnight 37°C). From these, single colonies were mixed with sterile distilled water to attain a standard optical density (10 MacFarland units), and 10 μl spread onto iso-sense plates and incubated overnight at 37 °C with Meropenem (10 μg), Ceftazidime (30 μg), Piperacillin/Tazobactam (75/10 μg), ciprofloxacin (5 μg), tobramycin (10 μg) antibiotic discs. Antibiotic susceptibility was determined by measuring the zone of inhibition and the results were interpreted as sensitive, intermediate resistant, according to EUCAST guidelines (16).

### Whole genome sequencing

For short-read sequencing, genomic DNA (500 ng) was extracted from overnight cultures using the DNeasy Blood and Tissue Kit (QIAGEN, Hilden, Germany) and mechanically fragmented for 40 s using a Covaris M220 (Covaris, Woburn MA, USA) with default settings. Fragmented DNA was transferred to a tube and library synthesis was performed with the Kapa Hyperprep kit (Kapa Biosystems, Wilmington MA, USA) according to the manufacturer’s instructions. TruSeq HT adapters (Illumina, SanDiego CA, USA) were used to barcode the libraries, which were each sequenced in 1/48 of an Illumina MiSeq 300 bp paired-end run at the Plateforme d’Analyses Génomiques of the Institut de Biologie Intégrative et des Systèmes (Laval University, Quebec, Canada).

For longer-read sequencing, genomic DNA (15 µg) was extracted from overnight cultures using a Promega Wizard Genomic DNA Purification Kit, quantified using a Qubit 3.0 fluoromiter (Qubit dsDNA broad range assay kit, Life Technologies) and tested for purity using a NanoDrop 1000 spectrophotometer (Thermo Scientific). Single Molecule Real-Time (SMRT) sequencing was performed at the Centre for Genomic Research, University of Liverpool. For each genome, three Single Molecule, Real-Time (RSII SMRT) cells with P6/C4 chemistry on a Pacific Biosciences (PacBio) were used to generate the raw sequence data.

### Genome assembly

For PacBio sequencing, the sequencing reads were *de novo* assembled using the HGAP v.3 workflow (17) through the Pacific Biosciences SMRT Portal v.2.3.0. The RS_HGAP_Assembly.3 protocol was run with default settings, except that the estimated genome size was set to 6.5 Mb. Contigs featuring either <20 x coverage, <20 Kb length, <47 Quiver-based quality values (QV) or > 95% sequence identity to larger contigs with better coverage and QV values were discarded from the assembly. Surviving contigs were assessed for circularization using the Circlator v.1.5.3 toolkit (18). Genes *dnaA* and *repA* were set as the start positions of the chromosomes and plasmids, respectively. Circularized contigs were further polished by following the RS_Resequencing.1 protocol through the SMRT Portal with default settings, and by mapping their corresponding Illumina-generated sequencing reads with BWA-MEM v.0.7.17-r1188 (arXiv:https://arxiv.org/abs/1303.3997v2) and correcting following the PILON pipeline v.1.22 (19).

PacBio genome assembly and genomes annotation was performed using a virtual machine hosted by the Cloud Infrastructure for Microbial Bioinformatics (CLIMB) consortium (20). For Illumina sequencing, raw Fastq files were processed for adapters and quality trimming using Cutadapt v.1.2.1 (21) and Sickle v.1.200 (https://github.com/najoshi/sickle) as previously reported (22). The genomes were *de novo* assembled and scaffolded from the filtered sequencing reads following the A5 MiSeq assembly pipeline (23) as described previously (22). Genome assemblies quality was assessed using QUAST v.3.1 (24).

### Annotation

The multi locus sequence type (MLST) profiles of the genomes were identified from the pubMLST *Pseudomonas aeruginosa* scheme (http://pubmlst.org/paeruginosa/)(25) using the mlst tool v.2.8 (https://github.com/tseemann/mlst).

AMR genes were detected in the megaplasmids sequences through BLASTn searches (26) against the Comprehensive Antibiotic Resistance Database (CARD) (27) using the ABRicate tool v.0.8 (https://github.com/tseemann/abricate). Only matches displaying coverage >30% were selected to generate the heatmap representing the AMR gene content of the megaplasmids group with the ComplexHeatmap package v.1.17.1 (28).

Genome annotation was carried out with Prokka v.1.12 (29) for both the Thai sequences reported in this study and the complete megaplasmids identified in GenBank (Table 1) to homogenize the annotation process. All the sequences were permuted to set the *repA* gene as the start position prior to annotation. Isolates genome annotation was complemented with the Sma3s tool v.2 (30) and by integrating data from conserved domains searches performed with InterProScan v.5.30-69.0 (31, 32). Integrons occuring in the Thai megaplasmids were identified using the Integron Finder package v.1.5.1 (33–36).

**Table 1.** Complete megaplasmids of the pBT2436-like family.

| Plasmid | Bacterial species | Strain | Size | Number of proteins | Country | Source | Accession Number | Reference |
| --- | --- | --- | --- | --- | --- | --- | --- | --- |
| pBT2436 | *P. aeruginosa* | 2436 | 422,811 | 536 | Thailand | Respiratory infection | CP039989 | This work |
| pBT2101 | *P. aeruginosa* | 2101 | 439,744 | 556 | Thailand | Respiratory infection | CP039991 | This work |
| unnamed2 | *P. aeruginosa* | AR_0356 | 438,531 | 556 |  |  | CP027170 |  |
| unnamed2 | *P. aeruginosa* | AR439 | 437,392 | 545 |  |  | CP029096 |  |
| unnamed3 | *P. aeruginosa* | AR441 | 438,529 | 556 |  |  | NZ_CP029094 |  |
| pJB37 | *P. aeruginosa* | FFUP_PS_37 | 464,804 | 597 | Portugal | Respiratory infection | KY494864 | Botelho et al. 2017 |
| pBM413 | *P. aeruginosa* | PA121617 | 423,017 | 542 | China | Respiratory infection | CP016215 | Liu et al. 2018 |
| pOZ176 | *P. aeruginosa* | PA96 | 500,839 | 623 | China | Respiratory infection | KC543497 | Xiaong et al. 2013 |
| p12939-OXA | *P. aeruginosa* |  | 496,436 | 608 | China* |  | MF344569 |  |
| p727-IMP | *P. aeruginosa* |  | 430,173 | 534 | China* |  | MF344568 |  |
| pA681-IMP | *P. aeruginosa* |  | 397,519 | 486 | China* |  | MF344570 |  |
| pR31014-IMP | *P. aeruginosa* |  | 374,000 | 456 | China* |  | MF344571 |  |
| pRBL16 | *P. citronellolis* | SJTE-3 | 370,338 | 485 | China | Wastewater sludge | CP015879 | Zheng et al. 2016 |
| p1 | *P. koreensis* | P19E3 | 467,568 | 599 | Switzerland | *Origanum marjorana* | CP027478 | Schmid et al. 2018 |
| pSY153-MDR | *P. putida* | SY153 | 468,170 | 580 | China | Urinary tract infection | KY883660 | Yuan et al. 2017 |

\* Country of isolation is putative, based on the information of the submitter's country in GenBank.

### Comparative analysis

Pairwise comparison of the pBT2436 and pBT2101 megaplasmid sequences was performed through BLASTn (26) and visualized with ACT v.17.0.1 (37). Repeated sequences in the pBT2436 and pBT2101 resistance regions were identified by aligning the regions against themselves with BLASTn and visualizing the output with Aretims Comparison Tool (ACT) masking the largest match corresponding to the overall sequence identity. Sequences homologous to the pBT2436 Resistance region 2 encoding the MexC-MexD-OprJ efflux pump were detected through BLASTn (26) searches against the Non-redundant GenBank database and the pairwise alignments visualized with ACT v.17.0.1 (37).

Complete pBT2436-like megaplasmid sequences deposited in GenBank were found through BLAST searches (26) against the Non-redundant nucleotide collection using pBT2436 as query sequence. The search was performed in November 2018 and only hits featuring <1e-5 E-value, >75% Query coverage and >1e5 Max score were selected as complete homologous sequences. Note that all the non-selected sequences displayed query coverage values below 25%. A circular representation of the genome comparison between the megaplasmids detected in GenBank and pBT2436 or pBT2101 at nucleotide level was generated using the BRIG application v.0.95 (26, 38, 39).

The pangeome analysis of the pBT2436-like megaplasmids group was performed with the GET_HOMOLOGUES package v.10092018 (40–47). The GET_HOMOLOGUES pipeline was ran with default settings except for the additional “-E 1e-03 -c -z” options. Core and Pan-genome were computed with the compare_clusters.pl script as the intersection of the BDBH-COG-OMLC and COG-OMCL algorithms, respectively. Rarefaction curves were generated using the plot_pancore_matrix.pl script. The pan-genome matrix obtained from the analysis was visualized as a heatmap with the ComplexHeatmap package v.1.17.1 (28). Functional annotation of the accessory proteins of the group was performed using Sma3s v.2 (30).

The phylogenomic analysis of the megaplasmids group was carried out following the GET_PHYLOMARKERS pipeline v.2.2.5_9May18 (48–57). The set of single-copy core-genome clusters detected using GET_HOMOLOGUES was input into the pipeline to identify high-quality phylogenetic markers and inferring a core genome phylogeny through a maximum-likelihood tree search. The pipeline was run under the following settings: phylogenetics mode, DNA sequences, IQ-TREE as the tree-searching algorithm, “high” model evaluation mode for IQ-TREE (based on ModelFinder) during the gene-trees searching, and 10 independent IQ-TREE tree searches on the concatenated top-scoring markers (“-R1 -t DNA -A I -T high -N 10” options). A pan-genome phylogeny was inferred from the gene content profiles of the members of the group using the estimate_pangenome_phylogenies.sh script. A maximum-likelihood tree search was performed on the pan-genome matrix built from the GET_HOMOLOGUES analysis using the binary and morphological models implemented in IQ-TREE and launching 10 independent tree searches. The phylogenetic trees were visualized and edited with the iTOL tool v.4.3.3 (58).

### Megaplasmids homologues search

The corrected Illumina-generated sequencing reads from the Thai isolates were mapped to the plasmid pBT2436 searching for related megaplasmids using BWA-MEM v.0.7.17-r1188 (arXiv:https://arxiv.org/abs/1303.3997v2). The presence of megaplasmids was assessed from the percentage of reads mapped to pBT2436 extracted from the alignment files with samtools v.1.7 (59). Genomes displaying less than 1% of mapped reads were considered as lacking the pBT2436-like megaplasmids based on the value obtained for the isolate 4068 genome used as negative control. Coverage of the pBT2436 sequence was assessed and visualized with BRIG v.0.95 (38, 39).

The search of megaplasmids homologous to pBT2436 in the larger database was performed with NUCmer from the MUMmer package v.3.1 (60). The query dataset was obtained from the GenBank entries reported at (61), and the collection of *Pseudomonas* genomes available at the GenBank assembly database (https://www.ncbi.nlm.nih.gov/assembly) in November 2018. The query sequences were aligned to pBT2436 running nucmer with default settings but using anchor matches that were unique to both the reference and query sequences (-mum). pBT2436 percentage of coverage was estimated by adding up the reference coverage values of all the alignments listed in the .coords files generated with the show-coords utility. Plots displaying both the pBT2436 coverage and percentage of sequence identity and location of the alignment regions were generated using the mummerplot script. Metadata associated to the queries matching the pBT2436 sequence were extracted from their corresponding biosample records.

### Accession numbers

The accession numbers of the Illumina-sequenced genomes from the Thai isolates are listed in Supplementary Table 1. The genome sequences of the 2101, 2436 and 4068 isolates generated with the PacBio platform have been filed under the GenBank BioProject PRJNA540594.

## Results

### Antimicrobial susceptibilities of clinical isolates

To provide a snapshot of circulating *P. aeruginosa* in a hospital in Thailand, we assembled a collection of *P. aeruginosa* associated with different kinds of infection. In order to identify MDR isolates, we carried out initial susceptibility tests on all 48 clinical isolates for five antibiotics representing different modes of action (Supplementary Table S1). Nine of the 48 isolates were resistant (either full or intermediate) to all five antibiotics, whilst a further six were resistant to four of the five antibiotics.

### Genome sequencing identified related MDR plasmids in two clinical isolates from Thailand

We selected three of the most resistant isolates (2101, 2436 and 4068) for long-read (PacBio) genome sequencing. Chromosomes of the strains 2436 and 2101 were assembled in complete circularised sequences of 6782092 and 6573638 bp, for which 6214 and 6041 protein-coding genes were predicted, respectively. The chromosome of the strain 4068 was assembled in four contigs accounting for 7.1 Mb of sequence information. All the chromosome sequences featured a GC content of ∼66%. In each of the three strains we identified circularised plasmid sequences. Isolates 2101 and 2436 carried related megaplasmids (named pBT2436 [423 kb] and pBT2101 [440 kb] respectively), and harbouring multiple AMR genes. Isolate 4068, which contained a much smaller plasmid (51 kb) with no identifiable AMR genes, was not analysed further in this study. A pairwise comparison of pBT2436 and pBT2101 indicated that these plasmids shared approximately 90% of their genome, differing mostly in the regions carrying AMR genes (Resistance Regions RR1 and RR2, Supplementary Figure 1). In addition, there was a 7.7 kb region (Variable Region VR1) present in pBT2101 but absent from pBT2436, which was not related to AMR but was also present in the chromosome of isolate 4068. The annotations of pBT2436 and pBT2101 identified shared regions relating to replication and partitioning (*repA, parAB*), DNA transfer (*traGBV, dnaG* and type IV pilus-related / type II secretion genes), heavy metal resistance (*terZABCDEF*), and chemotaxis (*che* genes). Additionally, several genes encoding radical

S-adenosyl-L-methionine (SAM) enzymes were present in both plasmids (Supplementary Table 2). Although the two plasmids harbored *mer* operons, these differed both in gene content and sequence identity at the nucleotide level (*merRTPCA* in pBT2436 and *merRTPFADE* in pBT2101, which contains additional but divergent *merRTP*, Figure 1).

**Figure 1.**
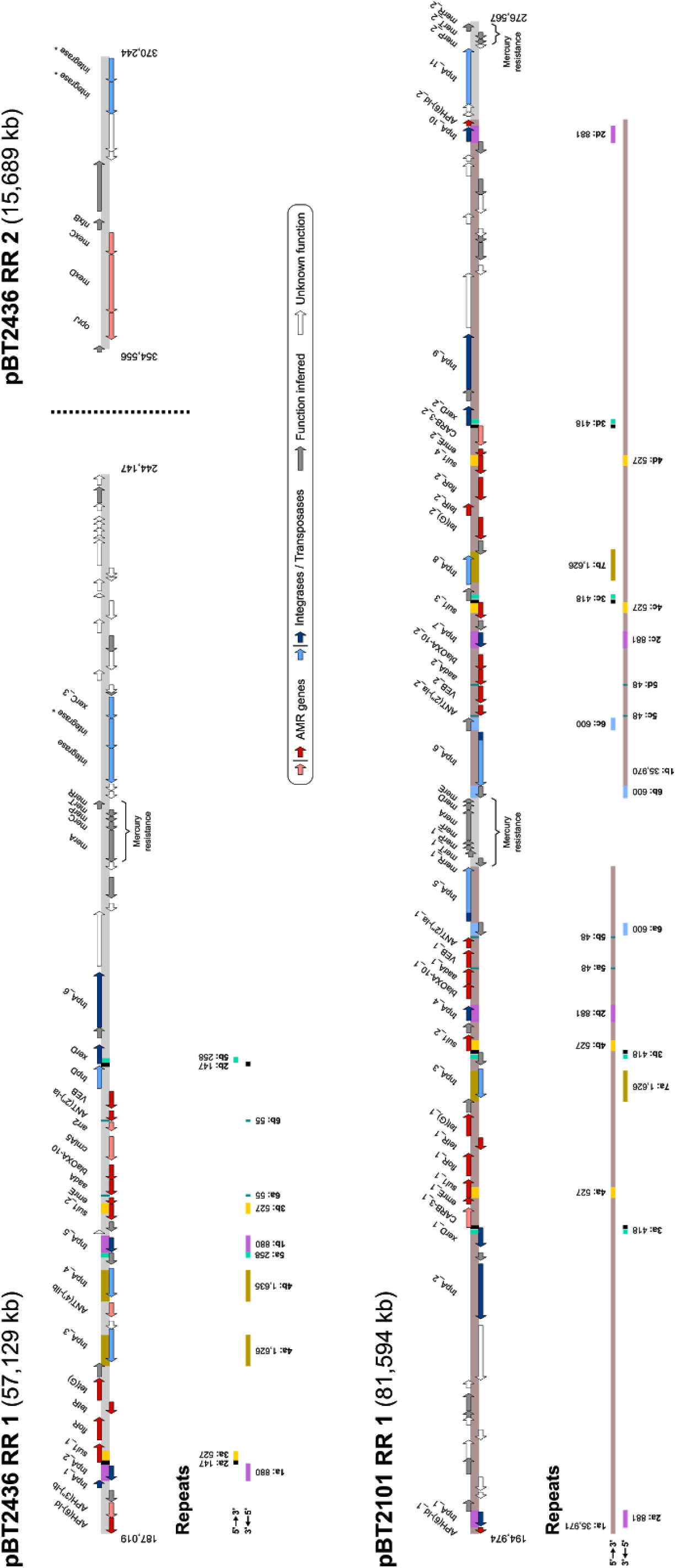
Maps of the pBT2436 and pBT2101 AMR regions. The coordinates of the AMR regions within the megaplasmids sequences are shown flanking the maps. ORFs and repeats, represented as coloured arrows and blocks, respectively, are indicated in the maps along with the location of the identified mercury resistance operons. Maps are drawn to scale. Gene names or products of selected ORFs of interest are indicated above their corresponding arrows. Gene duplications occurring within the same AMR region are numbered and distinguished with a “_N” suffix in their names. Darker shade-coloured arrows denote ORFs shared by the two megaplasmids’ RR1 regions. The different identified repeats are numbered according to their first occurence in the AMR region and a suffix letter (a to e) is included to distinguish between them. Repeats size in base pairs is displayed next to their corresponding blocks. Repeat blocks sharing matching colours between the two megaplasmids’ RR1 regions represent shared regions.

### AMR regions in pBT2436 and pBT2101 are mosaic and dynamic

Both plasmids carried an extensive array of AMR genes, encoding resistances against a wide range of antibiotics including beta-lactams (*VEB, blaOXA-10, CARB-3*), aminoglycosides (*ANT, APH* and *aad* genes), sulphonamides (*sul1*), tetracyclines (*tet* genes), macrolides (*ermE*) and phenicols (*floR*). We also identified genes for a chloramphenicol transporter (*cmlA5*), a ribosyltransferase (*arr-2*) and an efflux pump (*mexCD*-*oprJ*) present in pBT2436 (Figure 1).

Closer analysis of the AMR regions of the plasmids pBT2436 and pBT2101 revealed the presence of repeated sequences. In pBT2436, there was one large (RR1, 57.1 kb) and one small (RR2, 15.7 kb) Resistance Region containing AMR genes. In pBT2101 there was a single, large Resistance Region (RR1, 81.6 kb) (Figure 1). By aligning each of the two larger Resistance Regions with themselves, it was possible to identify both large and small duplicated regions, most notably a 35.9 kb region duplicated but inverted in pBT2101 when compared to pBT2436, with a unique central region of 4.36 kb in between the two duplicated areas containing a *mer* operon (Figure 1 and Supplementary Figure 2).

It is notable that the Resistance Regions were all rich in transposases and integrases, both unique and shared, in proximity to AMR genes, and there were a number of other interesting characteristics (Figure 1). There were regions shared between the two large Resistance Regions of pBT2436 and pBT2101. For example, they shared the *sul1-floR*-*tetR*-*tetG* region. However, in other areas there are partial matches where rearrangements have shuffled the order of genes. For example, in pBT2101 there was a cluster of resistance genes *blaOXA-10*-*aadA*-*VEB-1*-*ANT(2”)-Ia*, whereas in pBT2436, the genes were found in the order *VEB-2*-*ANT(2”)-Ia-arr2*-*cmlA5*-*blaOXA-10*-*aadA*. Likewise, the *emrE* gene occupied different positions in the two plasmids.

Within individual Resistance Regions there were also examples of both direct and inverted repeats (for example, *tnpA_3* and *tnpA_4*, and *sul1_1* and s*ul1_2* respectively in pBT2436), ranging in size from 48 to 1635 bp. We found that most of the repeats were shared between the Resistance Regions of the two plasmids but they differed in their arrangements (Figure 1). A further example of how dynamic these regions are can be seen with the repeats 2b and 5b of pBT2436, located next to *xerD* (Figure 1). Their pairs occupy different positions in pBT2436 RR1, whereas they occur merged as one repeat separated by 13 bp in the Resistance Region of pBT2101 (Figure 1).

We also saw examples of transposon insertions contributing to divergence between regions shared by the two megaplasmids. For example, the genes APH(6)-Id_1/2 in pBT2101, and *tnp*A_1 in pBT2436, represent truncated versions of their homologues due to the presence of adjacent *tnp*A_1/10 and *tnp*A_2 transposase genes, respectively (Figure 1).

In addition to predicted integrases, RR2 in plasmid pBT2436 carried genes for an efflux pump *mexC*-*mexD*-*oprJ* and a divergently transcribed *nfxB* gene, encoding the putative efflux pump repressor. Since this gene organization is identical to that present in the chromosome of the reference strain PAO1, we compared RR2 to the chromosome of the pBT2436-carrier strain to look for similarity at the nucleotide level (Supplementary Figure 3). Low levels of sequence identity (at best 78% sequence identity to a region comprising only *mexD*, and parts of *mexC* / *oprJ*), and differences in flanking genes suggested that the carrier strain chromosome was not the source of the RR2. Searches of the wider database revealed closer matches to sequences of diverse taxonomic origins, including both chromosomes and plasmids. The best match corresponded to a region in the chromosome of *Aeromonas hydrophila* strain WCHAH045096 (Accession CP028568), which harbors identity levels of 98% across the entire Resistance Region (Supplementary Figure 3). This region was found duplicated in the *A. hydrophila* chromosome (data not shown). Other close matches were found to regions of the non-related plasmids pBKPC18-1 (Accession CP022275) and pMKPA34-1 (Accession MH547560), from *Citrobacter freundii* and *P. aeruginosa* respectively (Supplementary Figure 3).The dynamic nature of these Resistance Regions, in contrast to the general conservation across the rest of the plasmid backbone, suggests that resistance genes have been assembled independently in the different plasmid backgrounds.

### Distribution of related megaplasmids amongst clinical isolates from a Thai hospital

In order to determine whether related plasmids were present in other *P. aeruginosa* from patients in the same hospital, a further 23 clinical isolates (including 2101 and 2436), chosen to represent a mixture of more resistant and less resistant isolates (Supplementary Table S1), were genome sequenced using the short-read Illumina platform. A summary of the genomics data obtained is shown in Supplementary Table S1. MLST profiles extracted from the genomes indicated that the sequenced isolates were highly diverse genetically.

In order to identify genomes harbouring plasmids related to pBT2101 and pBT2436, the Illumina reads were mapped on to the two plasmids as reference genomes. Four of the genomes (including those of the control strains, 2101 and 2436) mapped with coverage >90% (Supplementary Figure S4), indicating the presence of related plasmids in isolates 3582 and 638, both of which also displayed extensive AMR in our test (Supplementary Table S1). Isolates 2436 and 638, obtained from the same patient but from different types of sample (sputum and blood respectively), shared the same sequence type (ST-1121) and their plasmids shared high levels of identity. Isolates 2101 and 3582 also share the same sequence for six of the seven MLST loci but were isolated from different patients. Although the plasmids they carry share high levels of identity, when sequencing reads from isolate 3582 were mapped on to the pBT2101 genome, there was a clear reduction in numbers of reads mapping to the 81.6 kb Resistance Region containing the large duplication, consistent with the notion that the plasmid in isolate 3582 carries only one copy of the resistance genes rather than two (Supplementary Figure S4). Likewise, when 2101 and 3582 sequencing reads were aligned to the pBT2436 sequence, only 2101 displayed twice the number of reads mapping to the pBT2436 Resistance Region (Supplementary Figure S4).

Extended antimicrobial susceptibility tests were carried out on these four isolates, identifying some variations (Supplementary Table 3). For netilmicin, whereas for three of the isolates, zones of inhibition were close to the EUCAST breakpoint diameter of 12 mm (isolate 2101 just below and isolates 638 and 3583 just above), no zone of inhibition was detected for isolate 2436. When comparing the two strains carrying megaplasmids for which we obtained complete genomes, the only difference was that isolate 2101 was susceptible to amikacin.

### The family of pBT2436-like megaplasmids is widely distributed and present in non-*aeruginosa* species

Having identified the presence of members of the same family of megaplasmids (which we refer to as the pBT2436-like megaplasmids) associated with MDR in four of the clinical isolates from Thailand, we carried out homology searches targeting highly-similar complete sequences at the non-redundant NCBI nucleotide database. Details of the 13 additional megaplasmids identified, ranging in size from 370 kb to 501 kb and encoding from 456 to 623 proteins, are shown in Table 1.

Although ten of the megaplasmids were present in strains of *P. aeruginosa*, we identified related plasmids in three non-*aeruginosa* species of *Pseudomonas*, namely *P. putida, P. citronellolis* and *P. koreensis* (Table 1). The majority of isolates for which information about the country of isolation was available were linked with China and two of the three non-*aeruginosa* isolates were from sources other than human samples. The exception was the strain of *P. putida*, which was associated with a human urinary tract infection (62).

A comparative analysis of the 15 complete megaplasmids sequences revealed that they share high synteny and extensive sequence similarity, with the detected variation distributed at discrete loci across the genomes but mainly concentrated in large regions rich in AMR genes, transposases and integrases genes (Figure 2). The 7.7 kb variable region (VR1, Figure 2) previously detected in pBT2101 was absent from any of the other members of the megaplasmids group. Further homology searches focusing on VR1 only identified one match in GenBank, corresponding to the chromosome of the strain MRSN12280, which was recently described as the first report of a *P. aeruginosa* colistin-nonsusceptible isolate carrying the resistance gene *mcr* (63). However, annotation of the 7.7 kb pBT2101 unique region identified only some putative genes encoding transposable phage-related proteins, suggesting that this region does not contribute to the resistance. The GC content was similar among the members of the megaplasmids family (range 55.9-57.6%) although clear deviations from the average were detected mostly in the large variable regions (Figure 2). The GC content of the megaplasmids was lower than the median GC content reported for the genomes of their hosts, which range from 59.9% (*P. koreensis*) to 67% (*P. citronellolis*).

**Figure 2.**
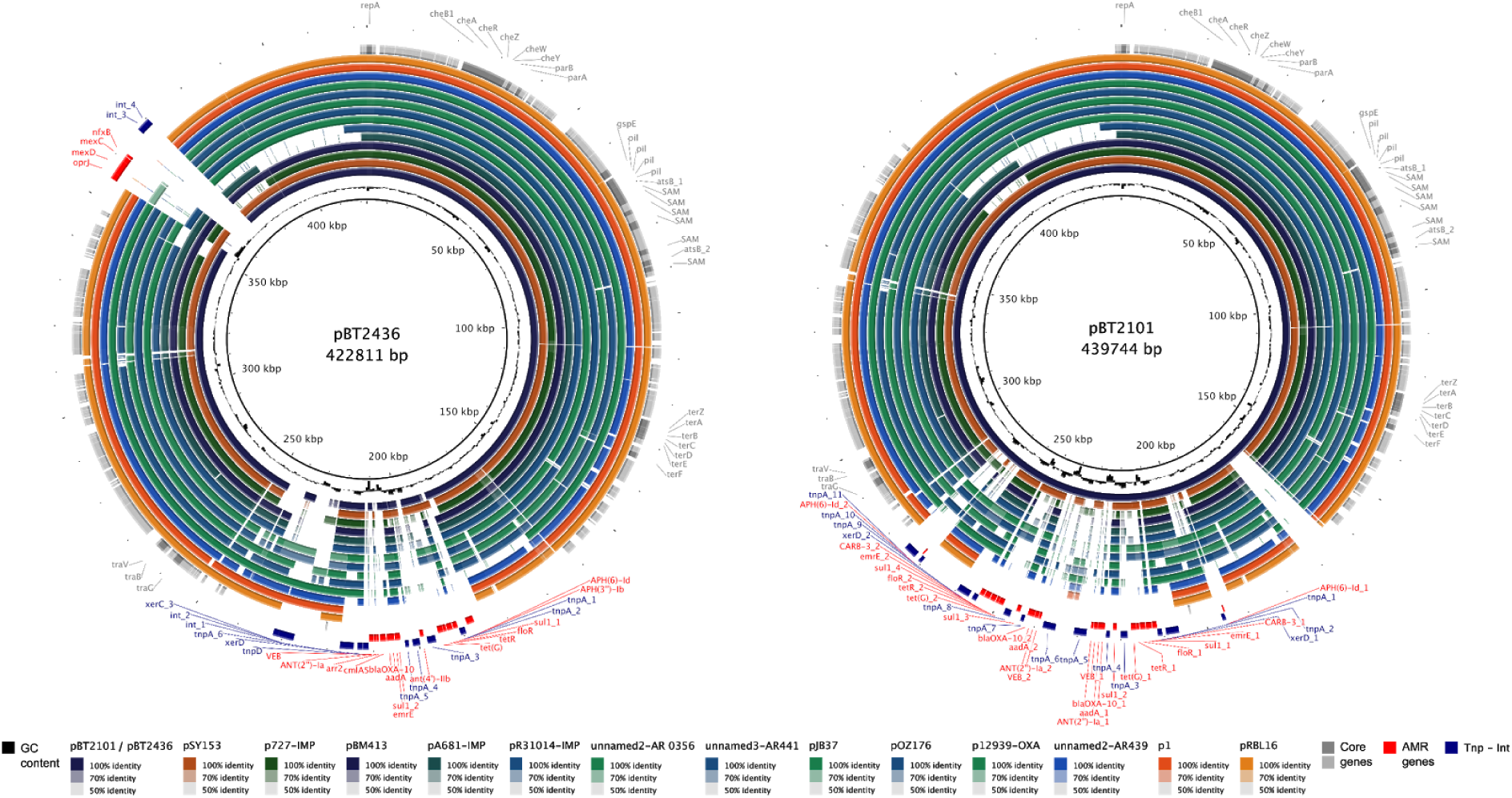
Genome comparison of megaplasmids of the pBT2436-like family. Fourteen complete *Pseudomonas* megaplasmids sequences, indicated at the bottom of the figure and represented as colour rings, were aligned to the pBT2436 (left) and pBT2101 (right) genomes at nucleotide level. Solid circles denote sequence homology to pBT2436/pBT2101 whereas gaps within the rings correspond to regions lacking of sequence similarity. Innermost rings (black) represent the GC content deviation from the average in the reference genomes. The three outermost rings (from innermost to outermost) indicate the location in pBT2436/pBT2101 of the core genes identified from the pan-genome analysis of the family (grey), AMR genes (red), and genes encoding integrases or transposases (blue). Names of genes of interest, and the position of the pBT2436/pBT2101 AMR (RR) and variable (VR) regions are indicated in the figure.

### Core genome and pan-genome analysis of the pBT2436-like megaplasmid family

Based on the comparative analysis of the 15 members of the megaplasmid family, we identified a core genome consisting of 261 orthologous protein groups, including proteins previously recognized as shared by plasmids pBT2101 and pBT2436 with roles in plasmid replication and partitioning, plasmid transfer, heavy metal resistance, chemotaxis, and a set of SAM proteins (Figure 2; Supplementary Table 2). On average, approximately 48% of each megaplasmid comprised the core genome (range 42-57%). Pangenome analysis using two different approaches yielded a consensus of 1164 orthologous protein groups (Supplementary Figure S5). Rarefaction and accumulation curves, used to estimate the core and pan-genome size of the megaplasmid family, were indicative of an open pan-genome with a well-defined core, and suggested that the addition of more genomes was unlikely to impact much further on the size of the core genome (Figure 3). Further analysis was carried out to distinguish between the strict core genome (present in all 15 megaplasmids), and other pan-genome compartments (40) (Figure 4, Supplementary Figure S5), indicating that the greatest number of gene clusters corresponded to unique, or nearly unique genes, but with numbers varying between the different megaplasmids from only one to 93 unique genes (Supplementary Figure S5, Supplementary Table 4), the latter being found in plasmid p1, a plasmid present in a *P. koreensis* isolate from healthy marjoram leaf material (64). Genes unique to plasmid p1 include a large cluster related to copper resistance and catabolic genes for the conversion of 4-hydroxy benzoate (*pobA, phbH*), or vanillate (*vanA, vanB, iacF, vanK*) to protocatechuate, which can then be metabolised to central metabolism intermediates via the β-ketoadipate pathway. Vanillate, a lignin derivative, and 4-hydroxybenzoate (also known as para-hydroxybenzoate) are both plant derived compounds.

**Figure 3.**
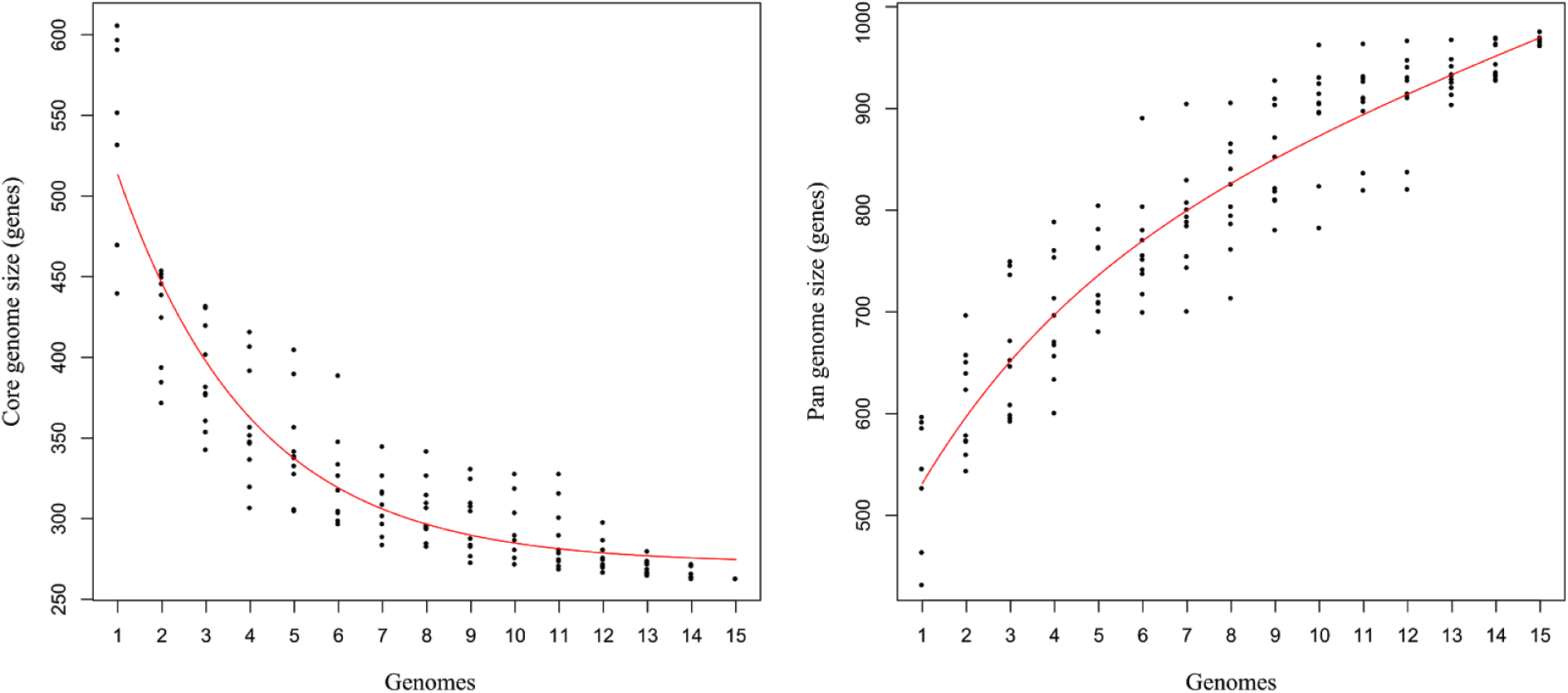
Rarefaction curves for the core and pan-genome of the pBT2436-like megaplasmids family. Graphs show the estimated size of the core (left) and pan-genome (right) of the pBT2436-like group. Plotted data comes from sampling experiments with ten random-seeded replicas from th BDBH-(40) (core) or OMLC-based (46) (pan) proteins clustering. Fitted curves follow the Tettelin function (47).

**Figure 4.**
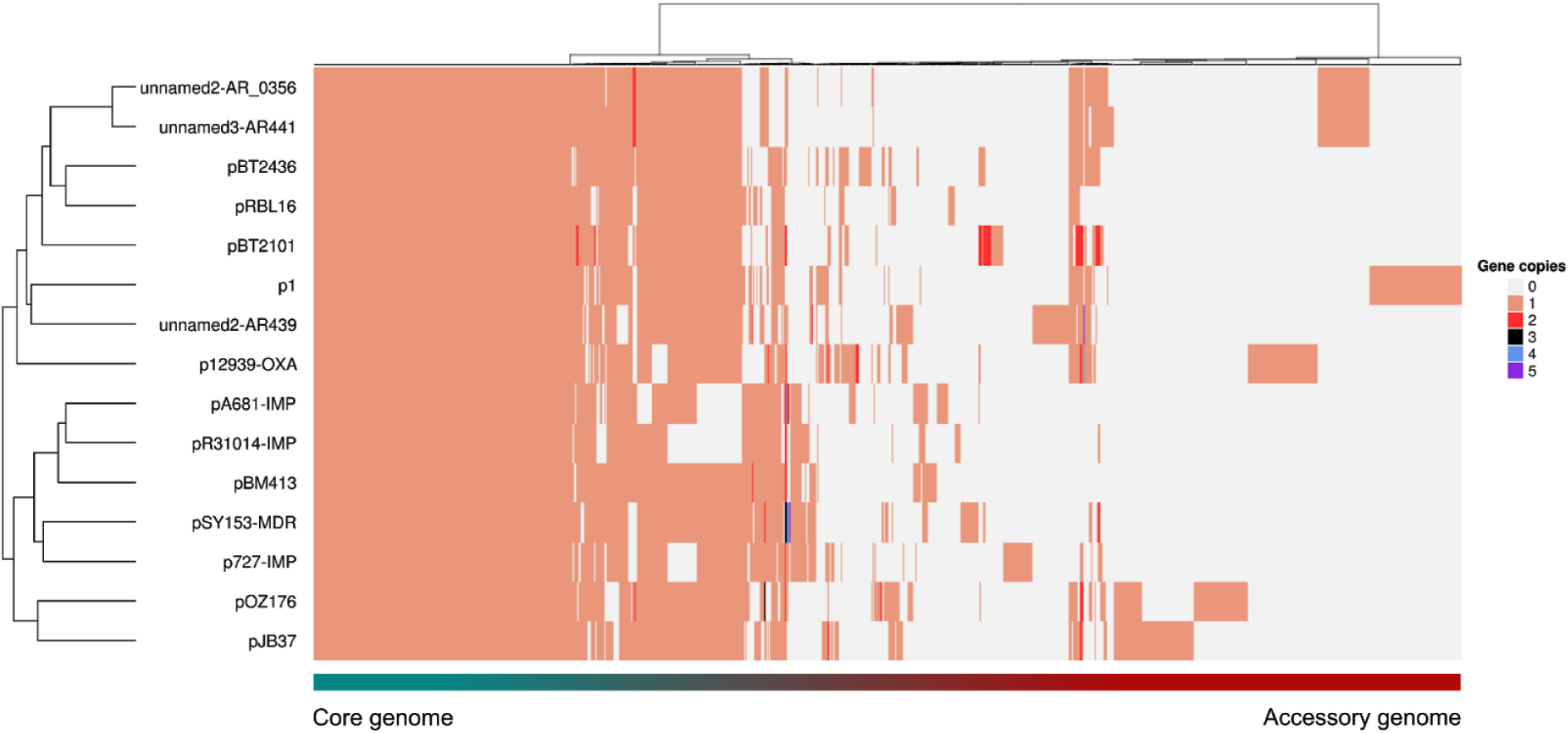
Pangenome matrix of the pBT2436-like megaplasmids family. The heatmap represents the gene content, in terms of presence/absence, of the pBT2436-like megaplasmids pan-genome inferred from the comparison of fifteen members with complete genomes. Comparison was performed at protein level rendering 1164 consensus groups as the result of the intersection between the COG-(41) and OMCL-based (46) clustering strategies (Supplementary Figure 5). Protein groups (X axis) and megaplasmids (Y axis) are hierarchically clustered based on the “ward.D” and “complete” methods, respectively, with Euclidean distance (28). Number of gene copies per cluster/per genome are colour-coded and numbered from 0 (grey:absence) to 5. Distribution of the pan-genome in the core and accessory components is indicated.

In the plasmids hosted by *P. aeruginosa* there were also clusters of “unique” genes, including a cluster of putative polysaccharide biosynthesis genes in the plasmid carried by *P. aeruginosa* AR439 and genes putatively involved in the oxidation of phosphite to phosphate in the plasmid p727-IMP. Other “unique” genes have putative functions related to antibiotic resistance, conjugation, transposases/integrases and chemotaxis, as well as many genes for which the role is not clear, including numerous with predicted regulatory functions. A breakdown of the functional categories predicted for the megaplasmid family accessory genomes is shown in the Supplementary Figure 6. Although functions were inferred for hundreds of accessory genes, the functions of approximately 70% of the megaplasmid family’s accessory component could only be characterised as “hypothetical proteins”. A similar proportion (approximately 73%) of genes with no indication of function was observed for the core component of the megaplasmid family’s pan-genome.

### Phylogenetic analysis and distribution of AMR genes among the pBT2436-like megaplasmid family

All the AMR genes identified in the megaplasmids were classed as part of the accessory genome, but several of them were carried on multiple plasmids. For example, the efflux pump genes detected in RR2 of pBT2436 were also present in six other members of the megaplasmid family (Figure 2). However, there was variation in flanking genes between the megaplasmids.

The pan-genome analysis identified several genes occurring in multiple copies (from 2 to 5) in members of the megaplasmids family (Figure 4). The majority of these genes were AMR genes. For example, the *P. putida* plasmid pSY153-MDR features four copies of the gene encoding the multidrug transporter EmrE, whereas pOZ176 harbours three copies of the gene *qacA* which is absent from the other megaplasmids and is involved in resistance against antiseptic and disinfectant compounds.

In order to better understand the relationships between the different members of the megaplasmid family, we constructed phylogenetic trees based on both the core-and the pan-genome (Figure 5). The core genome analysis was based on the nucleotide composition of a set of curated core genes (n = 105), whereas the pan-genome analysis was based on presence or absence of protein clusters. The two approaches yielded consistent clustering, in that we could identify a cluster of five megaplasmids isolated in China (including one found in *P. putida*), a cluster comprising the two megaplasmids pOZ176 (from China) and pJB37 (from Portugal), and a cluster of two megaplasmids for which we have very little information (from strains AR_0356 and AR441). The two other megaplasmids from non-*aeruginosa* hosts, and that from the *P. aeruginosa* strain AR439, do not cluster with any other plasmids, or each other. Interestingly, the two megaplasmids from the same hospital reported in this study (pBT2436 and pBT2101) cluster more closely by pangenome analysis than they do by core genome analysis, suggesting some convergence due to acquisition of accessory genes which, in these plasmids, are mostly AMR-related (Figure 5). When we carried out a comparative analysis of the AMR genes content in the megaplasmids family, the close relationship between the distribution of AMR genes carried by pBT2436 and pBT2101, albeit with variations, became apparent (Figure 6). In fact, the clustering seen in the phylogenetic trees was broadly reflected in the grouping obtained according to AMR gene content. It is also notable, that two of the megaplasmids found in non-*aeruginosa* strains (p1 and pRBL16), both from non-human sources, lacked any AMR genes, whereas plasmid pST153-MDR, isolated from a *P. putida* associated with a urinary tract infection in China, shares a similar AMR gene profile to other *P. aeruginosa* isolates from China (Figure 6).

**Figure 5.**
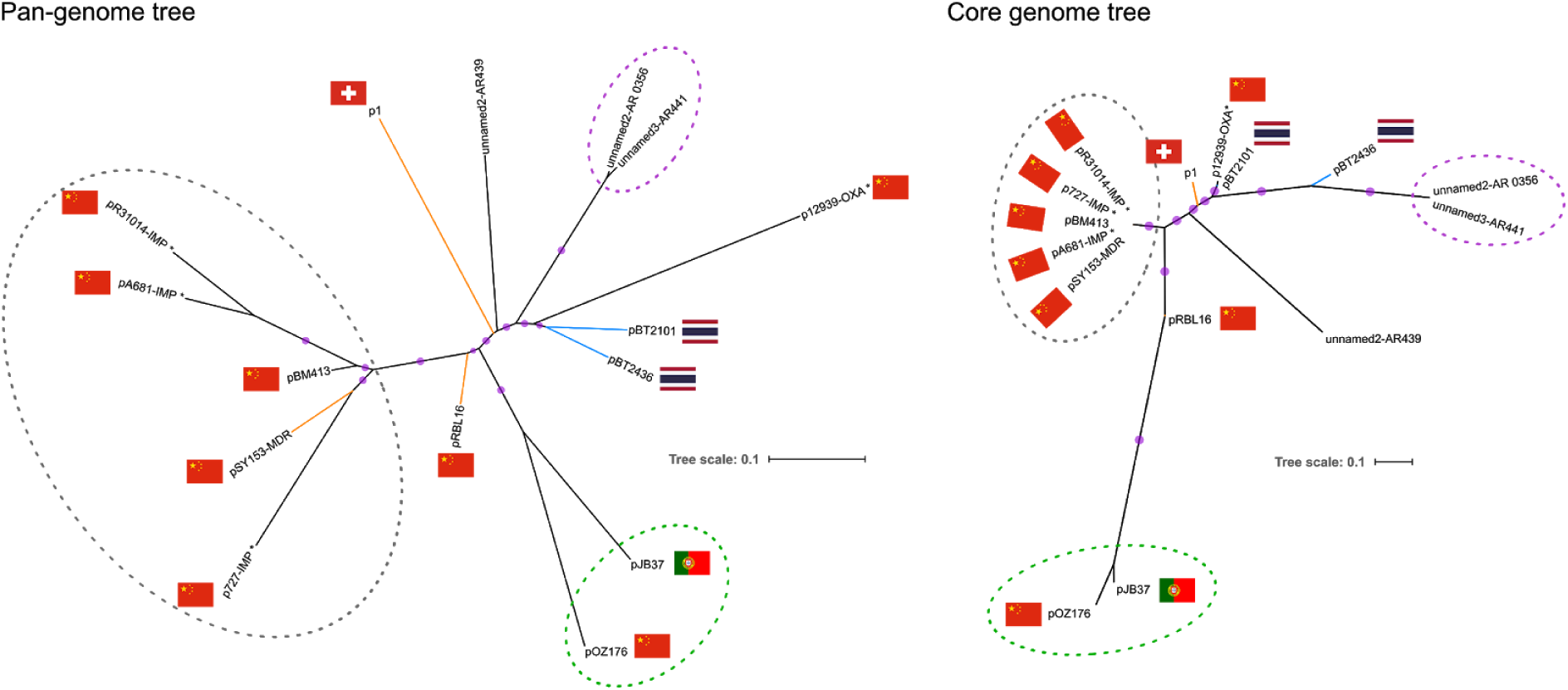
Phylogenomic analysis of complete megaplasmids of the pBT2436-like family. Maximum-likelihood unrooted phylogenetic trees displaying the relationships among the megaplasmids reported in this work (blue branches) and thirteen *Pseudomonas* homologous plasmids detected in GenBank. Branches corresponding to plasmids from non-*aeruginosa* species are coloured in orange. Pan-genome phylogeny (left) was estimated from the pan-genome matrix and reflects the evolutionary relationship of the megaplasmids in terms of their gene content, i.e. presence/absence patterns of the 1164 clusters contained in the matrix. The core genome phylogeny (right) was estimated from the concatenated set of 105 top-ranking alignments from core genes selected as phylogenetic markers and depicts the patterns of divergence of the megaplasmids genomic backbone. Taxa clustered together in both trees are highlighted with dotted-line ovals of matching colours. Purple dots represent approximate Bayesian posterior probability values > 0.99 and > 0.81 for the corresponding nodes in the core-and pan-genome trees, respectively. Scale bars indicate the number of expected substitutions per site under the binary GTR+FO (pan-genome tree) or best-fitting GTR+F+ASC+R2 (core genome tree) models. Where known, sources are indicated by nation flags. Asterisks in plasmid names denote cases where the geographical source is putative, based on the information of the submitter’s country in GenBank.

**Figure 6.**
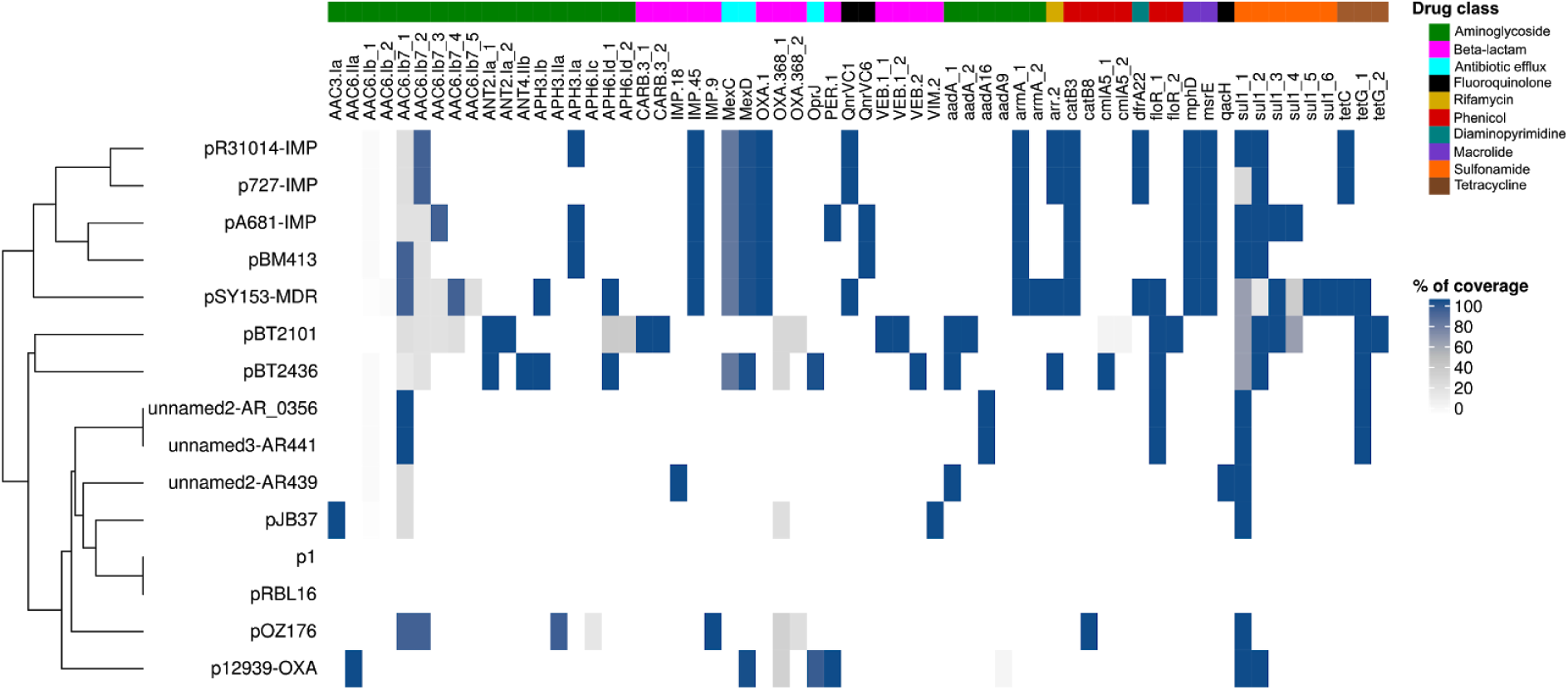
AMR genes content of the megaplasmids group pBT2436-like. The heatmap shows the collection of AMR genes identified in fifteen members of the pBT2436-like megaplasmids family trough BLASTn searches against the Comprehensive Antibiotic Resistance Database (CARD) (27). Percentage of coverage of the AMR genes identified from the searches is indicated. The megaplasmids (Y axis) are hierarchically clustered based on their content of AMR genes (X axis) using the “complete” method with Euclidean distance (28). The “_N” suffix in gene names denotes cases where the gene occurs more than once in the same megaplasmid. AMR genes are additionally classified based on the drug class they confer resistance to according to CARD (27).

### Wider distribution of the pBT2436-like megaplasmid family

We next sought to determine the wider distribution of the megaplasmid family by exploiting the rich source of genomic information available in public databases. We used a strategy of aligning nucleotide sequence data from genomes assembled to different levels, but mostly fragmented and produced by short-read sequencing technologies, to the complete pBT2436 nucleotide sequence.

Given the role of the megaplasmids in AMR, we first targeted a dataset of 390 genome sequences of *P. aeruginosa* clinical isolates from various infection and geographical sources and known to include many isolates associated with AMR (61). Our search led to the detection of ten genomes (∼2.6%) displaying estimated pBT2436 coverage values ranging from 21 to 95% (Supplementary Figure S7, Supplementary Table 5), with eight genomes having coverage values > 75%. These included genomes from isolates obtained from respiratory tract infections (4), intra-abdominal infections (2) and urinary tract infections (2), isolated in countries from Asia (China, India), the Americas (Mexico, Argentina and the United States) and Europe (Germany). All eight of these isolates were assigned to different MLST groups (61)(Supplementary Table 4). The oldest of these isolates was obtained in 2005.

Having established the approach, we targeted a wider database, searching for pBT2436-related megaplasmids in all the genomes from the *Pseudomonas* genus deposited in GenBank. At the time of our search more than 5,000 genomes were available at GenBank under different assembly categories for both *aeruginosa* (complete: 127; chromosome: 27; scaffold: 1053; contig: 1782) and *non-aeruginosa* (complete: 219; chromosome: 89; scaffold: 1266; contig: 1091) *Pseudomonas* species. Seventy-two matches (∼1.3%), 62 of them displaying >60% coverage values, were identified in this larger dataset (Supplementary Table 5). Most of the matches were detected from fragmented genomes (contig:33; scaffold:20). Metadata associated with the matching genomes was extracted from their biosample records (Supplementary Table 5). pBT2436-like megaplasmids were associated with strains from very diverse geographical origins (The Americas, Asia, Europe and Africa) and isolation sources, including both environmental and from a broad range of infection types (Figure 7, Supplementary Table 5). The best match, covering 96.6% of the pBT2436 sequence, was to a plasmid present in a strain of *P. montelii* isolated in Japan in 1986. The match with the oldest recorded collection date corresponded to a *Pseudomonas* sp. isolate from Japan dated back to 1970 and displaying coverage of 86.3% (Figure 7). Although most of the matches were identified in *P. aeruginosa* genomes, there were several examples of pBT2436-like megaplasmids found in other species of *Pseudomonas* (Figure 7, Supplementary Table 5)

**Figure 7.**
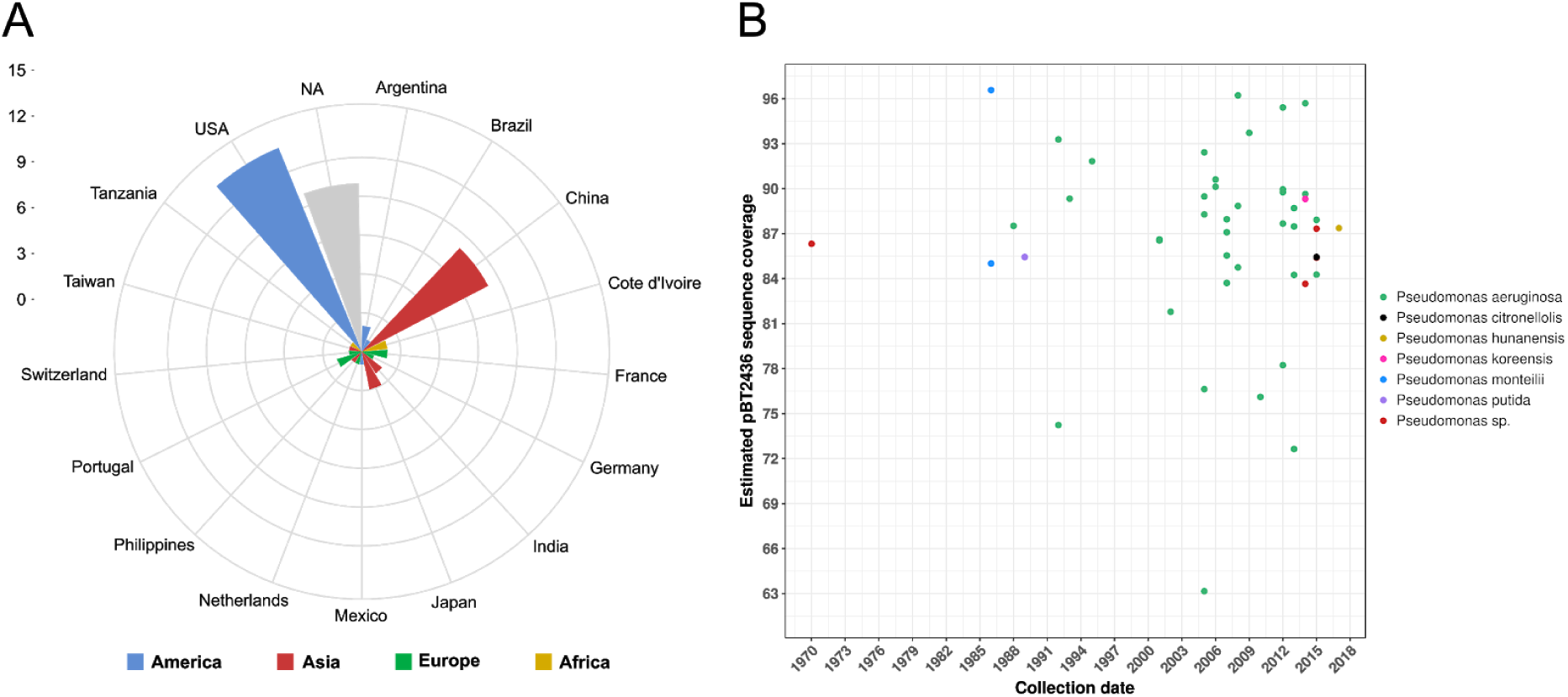
pBT2436-like megaplasmids recovered from Pseudomonas genome assemblies deposited in GenBank. Nucleotide sequences of *Pseudomonas* genomes from the four GenBank assembly categories were aligned to pBT2436 to identify unreported or overlooked related megaplasmids. Search was performed on November 2018. Plots show the geographic origin (A) and collection date (B) reported in the metadata (when available) of matches covering more than 60% of the pBT2436 sequence (Supplementary table 5). The bacterial species and pBT2436 estimated coverage from the matches is additionally indicated in B.

## Discussion

Surveillance is needed to monitor the emergence and spread of resistant clones of key pathogens such as *P. aeruginosa*, and whole genome sequencing approaches have now become sufficiently affordable that surveillance using such high-resolution technologies has entered the public health arena, especially in developed countries (65). However, the most accessible and affordable high-throughput sequencing approach relies on the use of short-read data, which can miss important information because of an inability to resolve repetitive or duplicated regions. Here we show the importance of generating complete genome data using a technology based on longer sequence reads. This enabled us not only to identify a family of MDR megaplasmids associated with the spread of resistance in a hospital in Thailand, but to identify subtle recombination and duplication events within two of the plasmids, indicative of mosaic AMR regions and dynamic evolution. The association of key adaptive traits with duplicated regions has also been reported for the p1 megaplasmid-carrying *Pseudomonas koreensis* strain P19E3 (64), with genes encoding heavy metal resistance, aromatic compound degradation, and DNA repair occurring within large repeats in the genome. Consistent with our observations, repetitive genes in p1 were identified close to transposase and phage genes suggesting a key role for mobile elements like transposons in shaping these large genome duplications (64).

The megaplasmids pBT2436 and pBT2101 reported in this work carry a diverse array of AMR genes. In comparison to pBT2101, pBT2436 carries two additional aminoglycoside resistance genes (APH(3“)-Ib, ANT(4’)-IIb), both of which have been previously linked to amikacin resistance (66, 67), the only antibiotic susceptibility difference we detected between the isolates 2436 and 2101, with isolate 2436 being resistant. Various mechanisms can contribute to amikacin resistance (68), including efflux pumps. Hence, it it also possible that the additional MexCD-OprJ efflux pump of pBT2436 contributes to amikacin resistance, though previous studies have suggested that MexCD-OprJ plays a lesser role than other efflux pumps in aminoglycoside resistance (69). Genes encoding MexCD-OprJ were also found in the chromosome of the carrier strain, isolate 2436. However, it is notable that the *mexCD-oprJ* region on plasmid pBT2436 displayed higher sequence similarity to chromosomes and plasmids from different organisms than to the chromosome of the carrier strain 2436, thus highlighting the capability of this megaplasmid family to collect and transport genetic information across a wide phylogenetic distance.

We were able to characterise the distribution of multiple repeated regions within and between the two plasmids pBT2436 and pBT2101, and identify the shuffling of genes, revealing high levels of dynamism in the megaplasmid AMR gene regions. Regions containing mercury resistance genes were also dynamic, with divergence in both gene content and sequence identity. The *mer* determinants in bacteria are commonly found on plasmids as part of transposons and display a wide variety of arrangements (70). Consistent with this observation the *mer* operons identified both in pBT2436 and pBT2101 were located next to transposases or integrases, although of different classes, implying distinct origins.

Our analysis of a small collection of clinical isolates from the same hospital in Thailand revealed four isolates carrying related megaplasmids, including two from isolates obtained from the same patient but from different clinical samples, and two from different patients but present in isolates sharing the same MLST (isolates 2101 and 3583). The latter is consistent with a cross infection or common source event, but it is notable that the plasmids have diverged with respect to a large duplication of AMR genes during the event. These isolates were collected in 2013. The presence of MDR (including carbapenem-resistance) megaplasmids in four of only 23 isolates analysed (∼17.4%) suggests that the megaplasmids may be playing an important role in the spread of resistance in this clinical setting, and that a more contemporary wider surveillance study is urgently needed.

After further exploration of the available repository of complete genomes, we identified several related plasmids, mostly harboured by strains from Asia (China), consistent with the notion of a family of carrier megaplasmids acting as vehicles for the spread of AMR genes. These included pBM413, a MDR megaplasmid (62, 71) carrying *qnrVC6* and *bla*_IMP-45_in *P. aeruginosa* strain Guangzhou-Pae617 (72), a sputum isolate from a patient suffering from respiratory disease, and pOZ176, containing two integrons harbouring *bla*_IMP-9_and *bla*_OXA-10_ respectively, and isolated from aMDR strain obtained from the sputum of an intensive care unit patient (PA96), following an outbreak of carbapenem resistant *P. aeruginosa* (73, 74). *Interestingly, the MDR megaplasmid pSY153-MDR, carrying bla*_IMP-45,_ was discovered in a *P. putida* isolate from the urine of a cerebral infarction patient in China (62). Despite being present in a different species, this plasmid is more closely related to the other plasmids isolated from China than to different members of the megaplasmid family, suggesting that between-species transmission occurred locally.

There were extensive variations in the carriage of AMR genes amongst members of the megaplasmid family, including two of the megaplasmids lacking AMR genes altogether, both isolated from non-*aeruginosa* species of *Pseudomonas*. The pRB16 megaplasmid was identified in *P. citronellolis* SJTE-3, an estrogen and polycyclic aromatic hydrocarbon degrading bacterium isolated from active sludge at a wastewater treatment plant in China (75). *P. koreensis* P19E3, harbouring the megaplasmid p1, was isolated from healthy marjoram (*Origanum marjorana*) leaf material during an isolation survey on an organic herb farm (Boppelsen, Switzerland) in 2014 (64). The presence on p1 of a cluster of genes encoding copper resistance suggests that whilst members of the megaplasmid family share a common backbone, they are flexible in the adaptive traits that they carry.

By mining databases dominated by contigs assembled from short-read data, we were able to gain valuable insights into the ecology and evolution of the megaplasmid family. Our findings indicated that members of the pBT2436-like megaplasmid family are widely distributed geographically, and have been isolated from various sources, both environmental (including sewage, sludge, high-arsenic soil, lake water, plant material and phenol treatment bioreactors) and clinical (including sputum, pneumonia, urine, feces, blood, keratitis, cerebrospinal fluid, and burn samples). In addition, we were able to detect plasmid sequences highly similar to pBT2436 in genomes from at least six different *Pseudomonas* species. Most of the matches corresponded to *P. aeruginosa,* which dominates in the databases. However, several matches from non-*aeruginosa* species displayed higher coverage values than others detected from *P. aeruginosa* isolates, suggesting active and recent megaplasmid transfer within the genus. This idea is consistent with the core and pan-genome trees constructed from complete megaplasmid sequences, where we observed a clear clustering of the *P. putida* megaplasmid pSY153-MDR with other members of the family hosted by *P. aeruginosa* isolates. Based on the serotype and MLST metadata available for some of the analyzed genomes, it is clear that the pBT2436-like megaplasmids are widely distributed within the *P. aeruginosa* population.

By analysing databases that include genome data from isolates obtained decades ago, we were able to obtain some insights into the temporal development of the megaplasmid family. Although most of the isolates where we detected matches were isolated after the year 2000, reflecting the fact that the database is dominated by more recent isolates, nine of them were collected prior to the year 1995. The three oldest isolates, dating back to 1970-1986, corresponded to non-*aeruginosa Pseudomonas* species from Japan. It is tempting to speculate on an evolutionary process whereby megaplasmids “empty” of AMR genes assembled in non-*aeruginosa* species, some collecting diverse cassettes of AMR genes either before or after moving into *P.aeruginosa*, with subsequent selection for the MDR-associated arrangements now seen in a clinical setting. Although much of the data that we present is consistent with this view, the best database match of all to our recent MDR megaplasmid pBT2436 (96.6% coverage), was to a plasmid present in a strain of *P. monteilii* isolated in Japan in 1986, suggesting that the reality is more complex, and that some plasmid/AMR arrangements that we might assume to be recently assembled, have existed for some time. However, it worth noting that *P. monteilii* has generally been associated with human pulmonary infections (76, 77) and that the 1986 isolate was isolated from a human source.

Further clues as to the origin and evolution of this megaplasmid family can be found through comparisons with more distant matches. Two environmental mercury resistance plasmids, pQBR103 and pQBR57, have similar gene order and content amongst backbone genes (78), including the presence of a radical SAM gene cluster, chemotaxis genes, and a type IV pilus, though they are both highly divergent at the nucleic acid level and thus were not detected in our survey. The conservation of this long region of synteny between distantly related plasmids suggests a functional association between the genes involved, but exactly what this is remains unclear. Neither pQBR57 nor pQBR103 contain any identifiable AMR genes and were isolated from pristine agricultural fields, consistent with the hypothesis of a pool of environmental megaplasmids, some of which have adapted to become vectors of clinically-relevant AMR.

Plasmid carriage is thought to impose fitness costs to host bacteria through the burdens associated with acquisition of superfluous genes, and for these megaplasmids, that increase the number of open reading frames by as much as ∼9%, these costs might be restrictive. However, discrepancy in size between large and small plasmids is also often balanced by reciprocal variation in copy number such that the total burden in terms of DNA replication is equivalent (79), and the contribution of size per se to plasmid cost is likely to be relatively small compared with costs associated with gene expression (80, 81). Megaplasmids may, therefore, be no less effective at AMR maintenance and transmission than smaller vectors. Furthermore, the fact that mutations can ameliorate even costly megaplasmids suggests that costs may not inhibit megaplasmid success in the long term (80, 82, 83). Our study highlights a widespread megaplasmid family present across clinical, environmental and geographical sources, and in multiple species hosts, suggesting widespread maintenance of this large plasmid backbone, and selection for maintenance of this backbone beyond adaptive traits such as AMR. The data provide important insights into the links between plasmids in the environment, acting as a reservoir for the emergence of important traits such as AMR, and MDR plasmids in a clinical setting.

## Author Contributions

AC and CW designed, conceptualized the study, and wrote the paper with input from the other authors. AC performed the bioinformatics analyses, the data curation and visualization. CW and JLF supervised the work and reviewed the manuscript. JLF participated in the study design. MPM, J-GE-R and RCL generated and assembled the Illumina-sequenced genomes. MG and LW performed the antimicrobial susceptibility testing. LW participated on isolating the genomic DNA. PP and PS collected the clinical isolates and the associated metadata.

## Acknowledgments

We would like to thank James P. J. Hall and Michael A. Brockhurst for their contributions after kindly reviewing this manuscript. This work was supported by Cystic Fibrosis Canada [R.C.L.], and the Secretaría de Educación, Ciencia, Tecnología e Innovación (SECTEI) [A.C.].

