## Supplemental Figures for "A megaplasmid family responsible for dissemination of multidrug resistance in *Pseudomonas*"

**Figure S1. Thai megaplasmids maps and pairwise comparison.** Left: pBT2436 and pBT2401 maps showcasing the location of AMR genes (red blocks) and genes encoding transposases or integrases (blue blocks). Innermost circle shows the GC content distribution of the megaplasmid genomes whereas the two outermost grey rings represent the ORFs encoded in the negative and positive strands. Right: Pairwise comparison of the pBT2436 and pBT2101 nucleotide sequences. Location of the Resistance (RR) and Variable (VR) regions of the megaplasmids is indicated.

**Figure S2. Duplications detected in pBT2436 and pBT2101 AMR regions.** Self-comparison of RR1 regions of the indicated megaplasmids at nucleotide level. Colour blocks connecting the AMR regions represent matches featuring >97% sequence identity either in the forward (red) or reverse complement (blue) direction. Overall self-similarity was masked to visualize discrete repeated regions. Note that the coordinates indicated correspond to the RR1 regions and not to the whole megaplasmid sequences.

**Figure S3. Comparative analysis of the pBT2436 Resistance Region 2.** The figure shows the pairwise comparisons at nucleotide level of sequences homologous to the pBT2436 RR2 (labeled in red). Compared regions correspond to chromosomal segments of the *P. aeruginosa* strains PAO1 and 2436 (pBT2436 carrier), the *A. hydrophila* strain WCHAH045096, and regions from the plasmids pBKPC18-1 and pMKPA34-1 of *C. freundii* and *P. aeruginosa*, respectively. The percentage of sequence identity detected in the homologous regions, depicted as green (forward) or magenta (reverse complement) connecting blocks, is indicated next to the corresponding matches. Coordinates of the compared regions in their corresponding genomes are shown flanking the maps, (c) denotes the sequence was changed to the reverse complement direction to easy the comparison visualization. ORFs of the regions are depicted as arrows and their colour code denotes: genes encoding the structural components of the MexCD-OprJ efflux pump (red) and its regulator NfxB (dark grey), genes encoding integrases (blue), and other genes with known (light grey) or unknown (white) function. Names of genes of interest are indicated above the corresponding arrows.

**Figure S4. Identification of pBT2436-like megaplasmids from short-read sequencing data of Thai *P. aeruginosa* clinical isolates.** The percentage of sequencing reads of the analyzed genomes mapping pBT2436 (green triangles) or pBT2101 (red dots) is plotted in A. Note that genomes of the strains 2436/2101 and 4068 represent positive and negative controls, respectively, as the presence/absence of pBT436-like megaplasmids in them was previously determined by long-read sequencing. The visualization of the reads alignment against the reference genomes for selected strains is shown in B. The first four rings (from the innermost to outermost) correspond to strains displaying the highest percentage of mapped reads whereas the next two rings represent the distribution of mapped reads of the negative control (4068, pink) and a strain categorized as lacking pBT2436-like megaplasmids (3979, grey). Blue regions in the rings denote mapping coverage values above the threshold and feature around twice the number of mapped reads than in the rest of the genome. Location of AMR genes, and those encoding integrases or transposases in the reference genomes, is shown in the two outermost black and purple rings, respectively . Resistance (RR) and variable (VR) regions in pBT2436 and pBT2101 are also indicated.

**Figure S5. Pangenome analysis of the pBT2436-like megaplasmid family.** A. Venn diagrams highlighting the intersection between the number of gene families computed by different clustering algorithms to identify the core (left) and pan-genome of the pBT2436-like megaplasmid group. BDBH: Bidirectional BLAST hits, COG: Cluster of Orthologous Groups triangle algorithm, OMCL: Orthologues Markov Cluster algorithm. B. Pan-genome structure analysis displaying the classification of the pan-genome consensus clusters into cloud, shell, soft-core and core compartments. C. Breaking-up of the number of sequence clusters identified per genome / per pan-genome compartment.

**Figure S6. Functional annotation of the pBT2436-like megaplasmids family accessory genome.** Protein annotation was performed with the Sma3s tool (30) and functional classification is based on the identified GO-terms (A and B) and Uniprot keywords (C) (Supplementary Table 4). Plots A and B show the number of accessory proteins classified into different "Molecular Function" and "Biological Process" GO-terms categories, respectively. The three plots compare the classification of all the accessory proteins of the megaplasmids group regarding those that were identified as unique in the pangenome, i.e. plasmid-specific proteins.

**Figure S7. Search of pBT2436-like megaplasmids in *Pseudomonas* genomes from the GenBank Assembly database.** Presence of pBT2436-like megaplasmids was assessed by aligning the query genomes to the pBT2436 nucleotide sequence and estimating its coverage from the alignments. Graph in A correspond to our pilot test and shows the pBT2436 coverage estimated from the alignments against a set of Illumina-generated genome assemblies of *P. aeruginosa* clinical isolates from Thailand. Positive and negative controls, where the presence or absence of pBT2436-related megaplasmids was previously determined by long-read sequencing or mapping of short-sequencing reads, are indicated. A visualization of the pBT2436 coverage (X axis) from the top-4 alignments, and their corresponding percentage of similarity (Y axis) is shown in B. Plots in C and D display similar data to A and B for the analysis of the 390 *P. aeruginosa* genomes reported by Kos et al. 2015.

### Figure S1. Thai megaplasmids maps and pairwise comparison

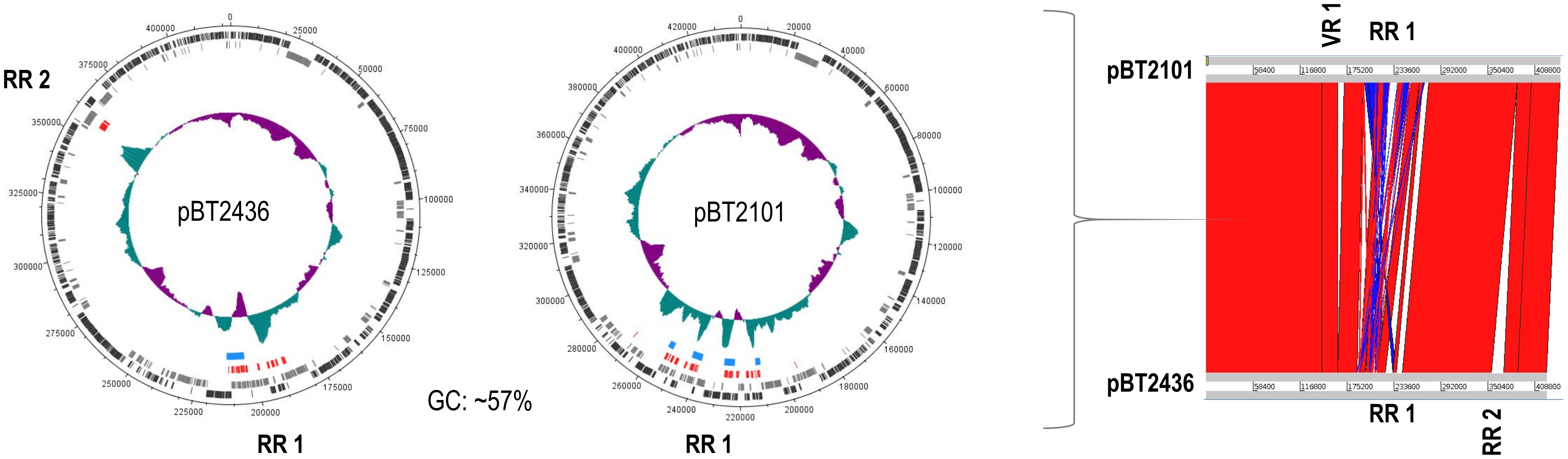

### Figure S2. Duplications detected in pBT2436 and pBT2101 AMR regions

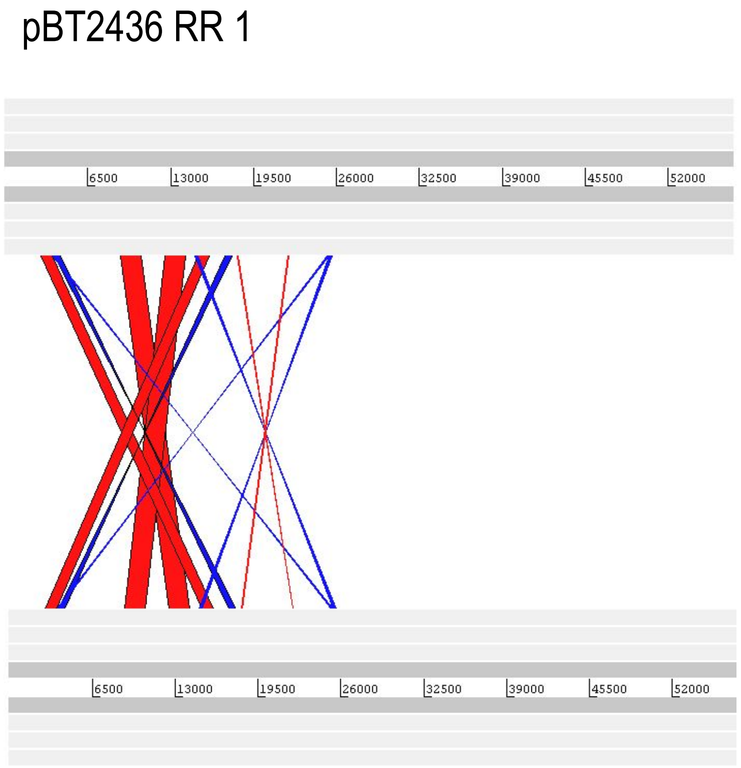


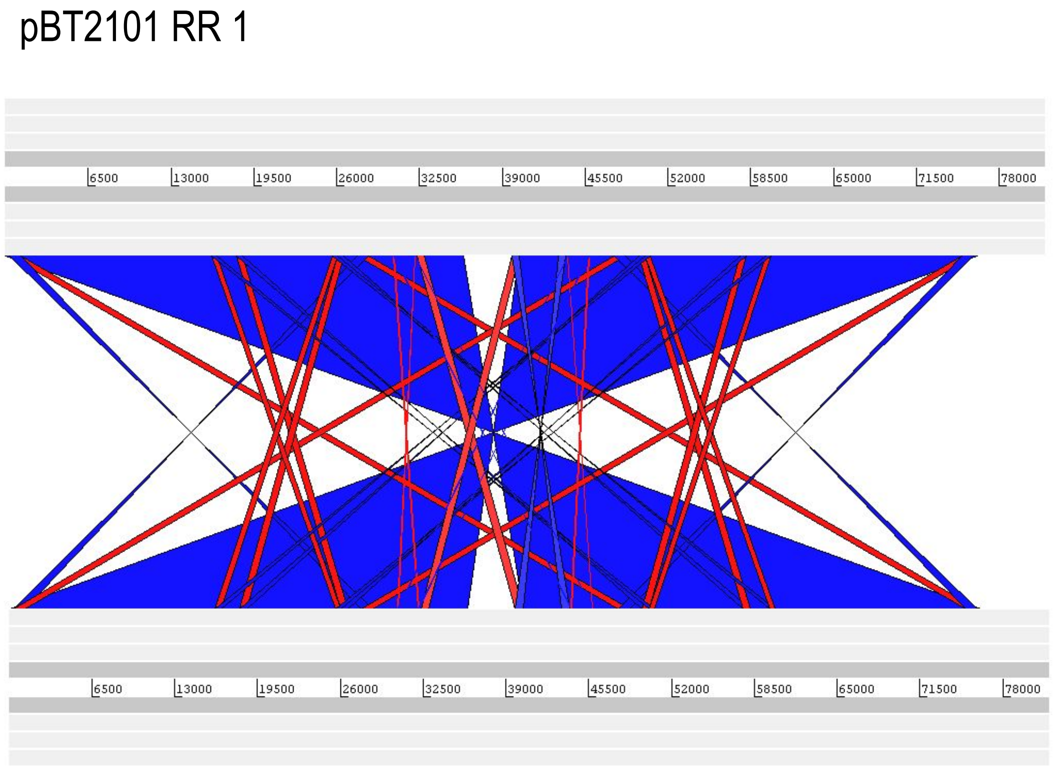

### Figure S3. Comparative analysis of the pBT2436 Resistance Region 2

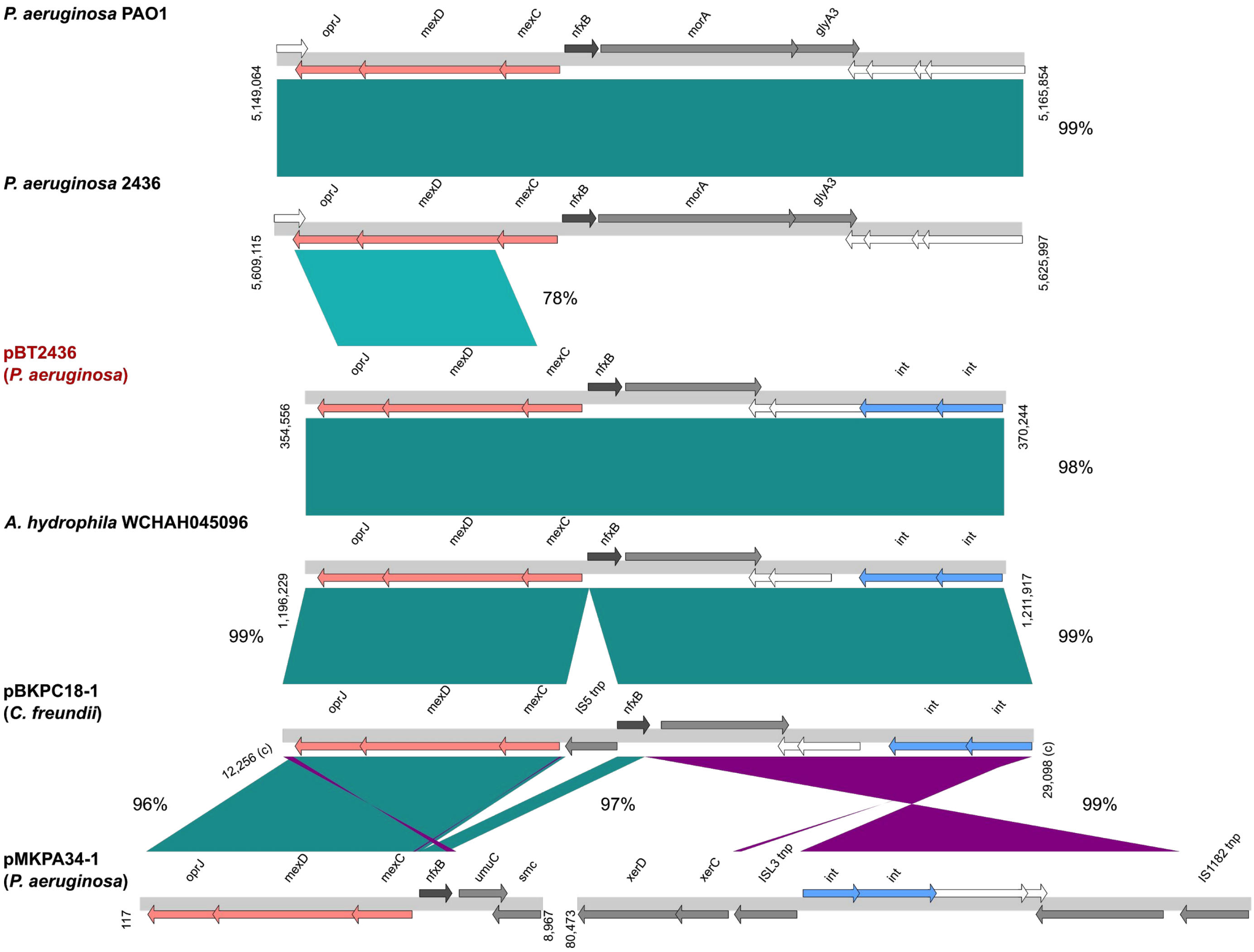

Figure S4. Identification of pBT2436-like megaplasmids from short-read sequencing data of Thai *P. aeruginosa* clinical isolates

**A**

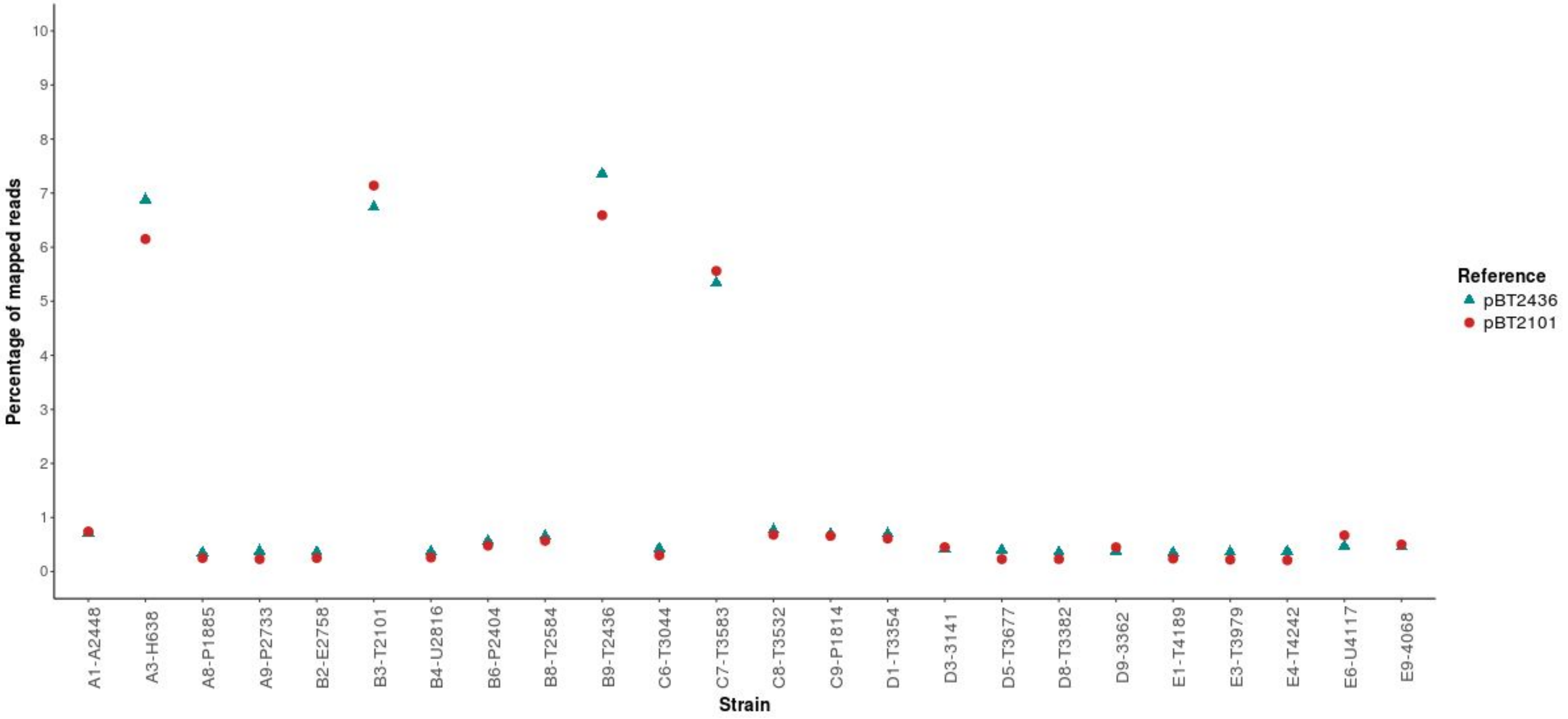

Figure S4. Identification of pBT2436-like megaplasmids from short-read sequencing data of Thai *P. aeruginosa* clinical isolates

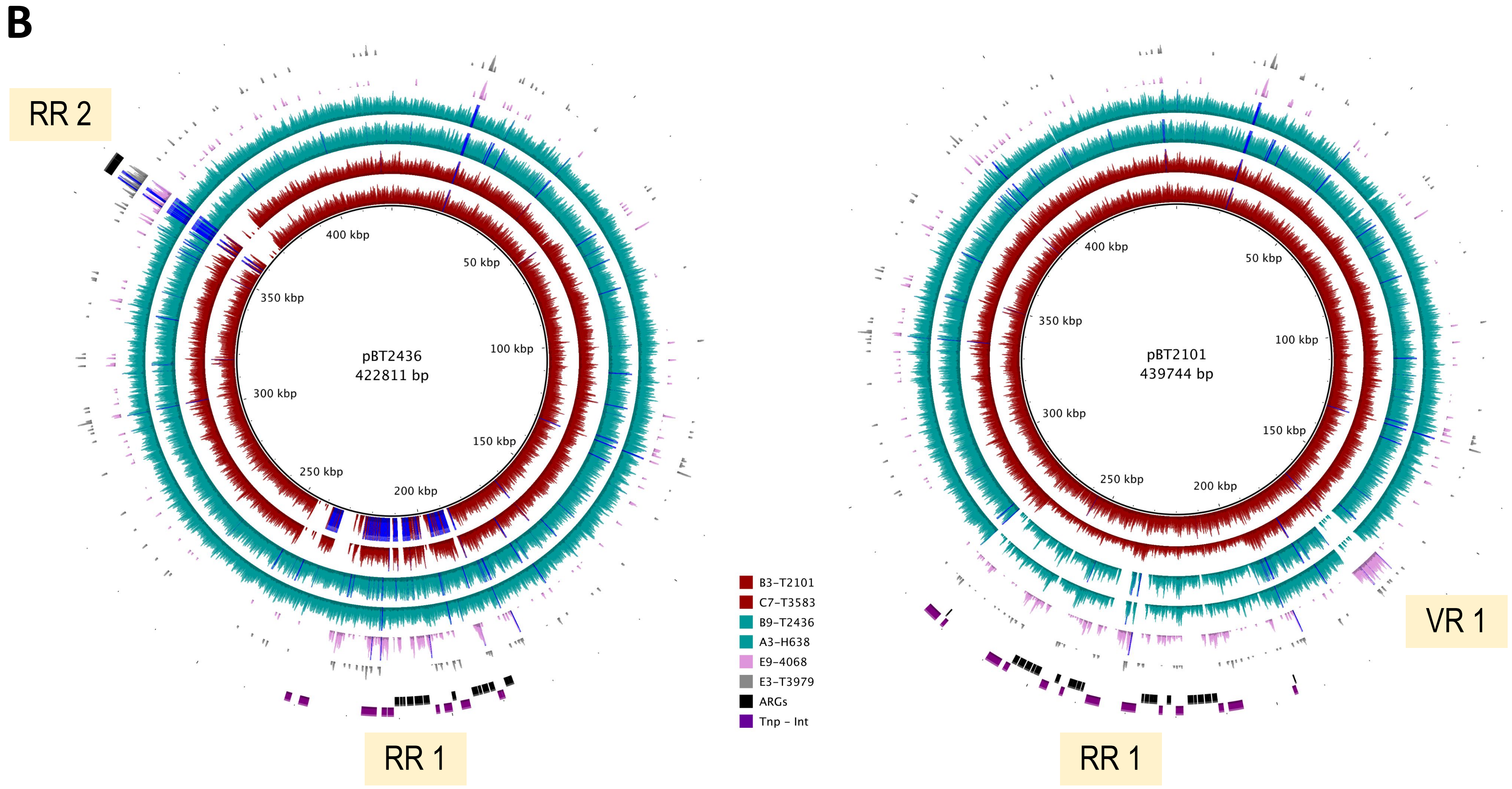

### Figure S5. Pangenome analysis of the pBT2436-like megaplasmid family

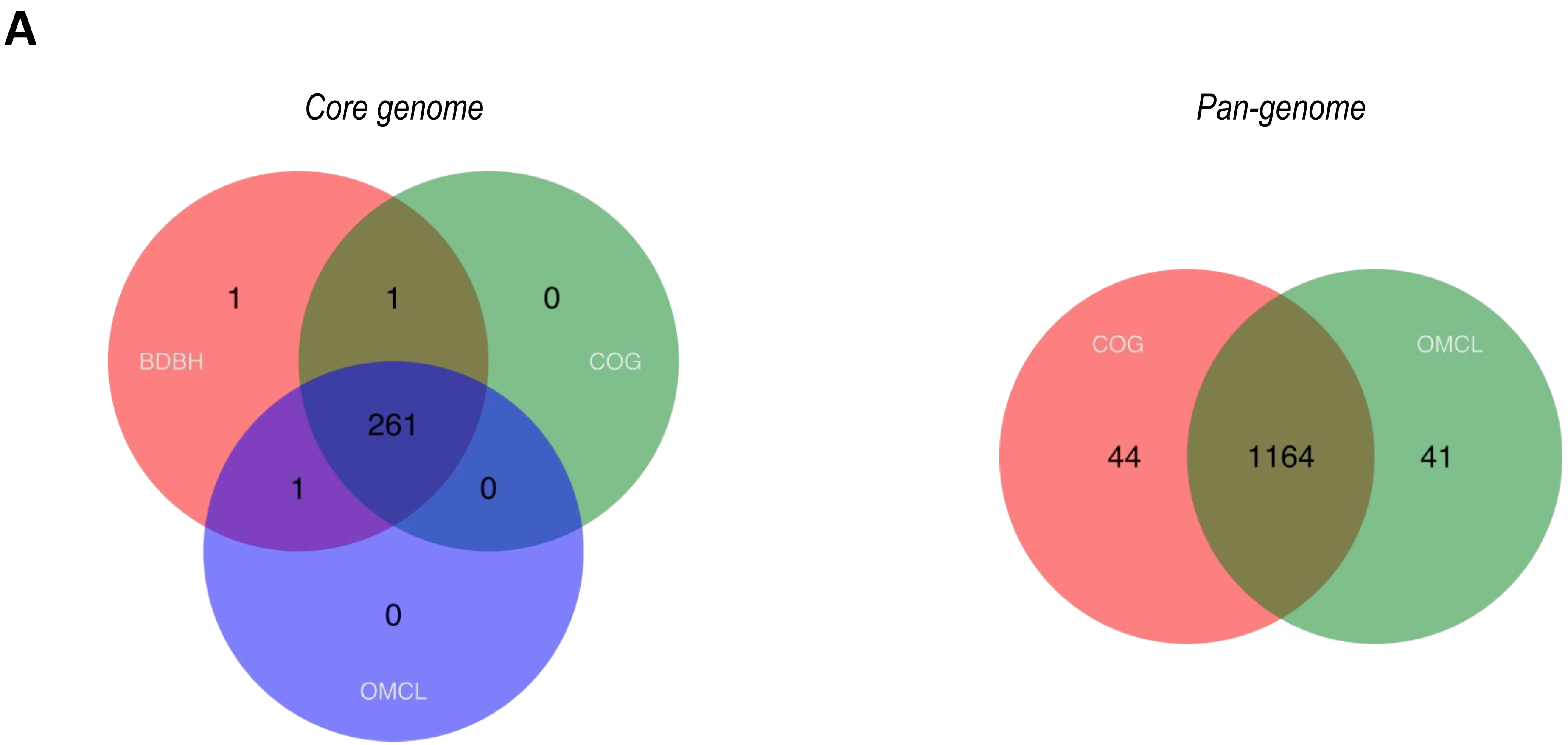


BDBH: Bidirectional BLAST hits
COG: Cluster of Orthologous Groups triangle algorithm
OMCL: Orthologues Markov Cluster algorithm

### Figure S5. Pangenome analysis of the pBT2436-like megaplasmid family

**B**

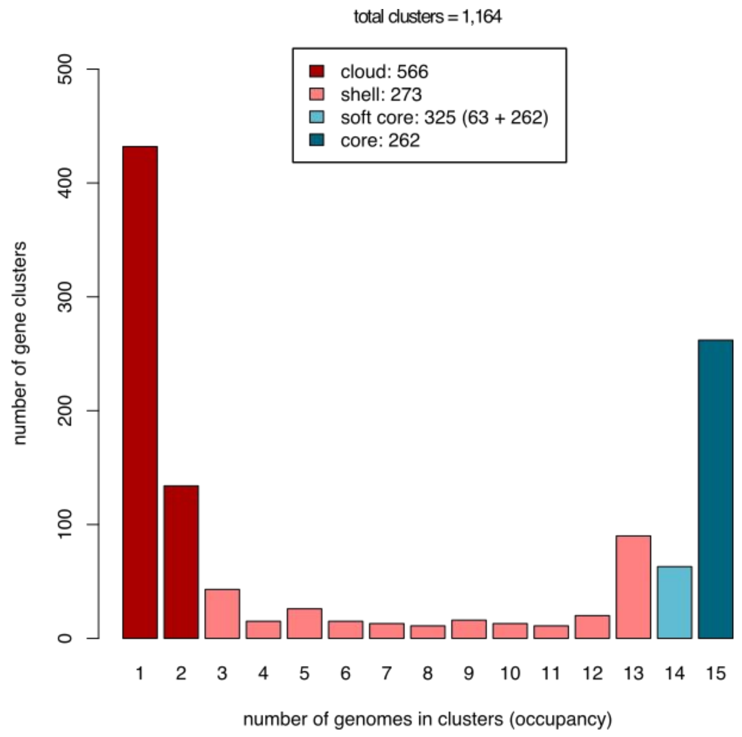


**C**

| Plasmid | Soft-core | Shell | Cloud | Unique |
|---|---|---|---|---|
| p1 | 324 | 168 | 98 | 93 |
| p12939-OXA | 308 | 169 | 96 | 75 |
| p727-IMP | 322 | 149 | 36 | 30 |
| pA681-IMP | 311 | 127 | 22 | 11 |
| pBM413 | 324 | 185 | 11 | 9 |
| pBT2101 | 324 | 161 | 22 | 18 |
| pBT2436 | 324 | 186 | 17 | 12 |
| pJB37 | 323 | 165 | 90 | 52 |
| pOZ176 | 317 | 169 | 103 | 57 |
| pR31014-IMP | 323 | 97 | 9 | 6 |
| pRBL16 | 323 | 151 | 9 | 7 |
| pSY153-MDR | 323 | 183 | 25 | 19 |
| unnamed2-AR439 | 316 | 159 | 50 | 37 |
| unnamed2-AR_0356 | 325 | 164 | 54 | 1 |
| unnamed3-AR441 | 325 | 160 | 58 | 5 |
| **Average** | | | | |
| | 320.8 | 159.5 | 46.7 | 28.8 |

### Figure S6. Functional annotation of the pBT2436-like megaplasmids family accessory genome

**A**

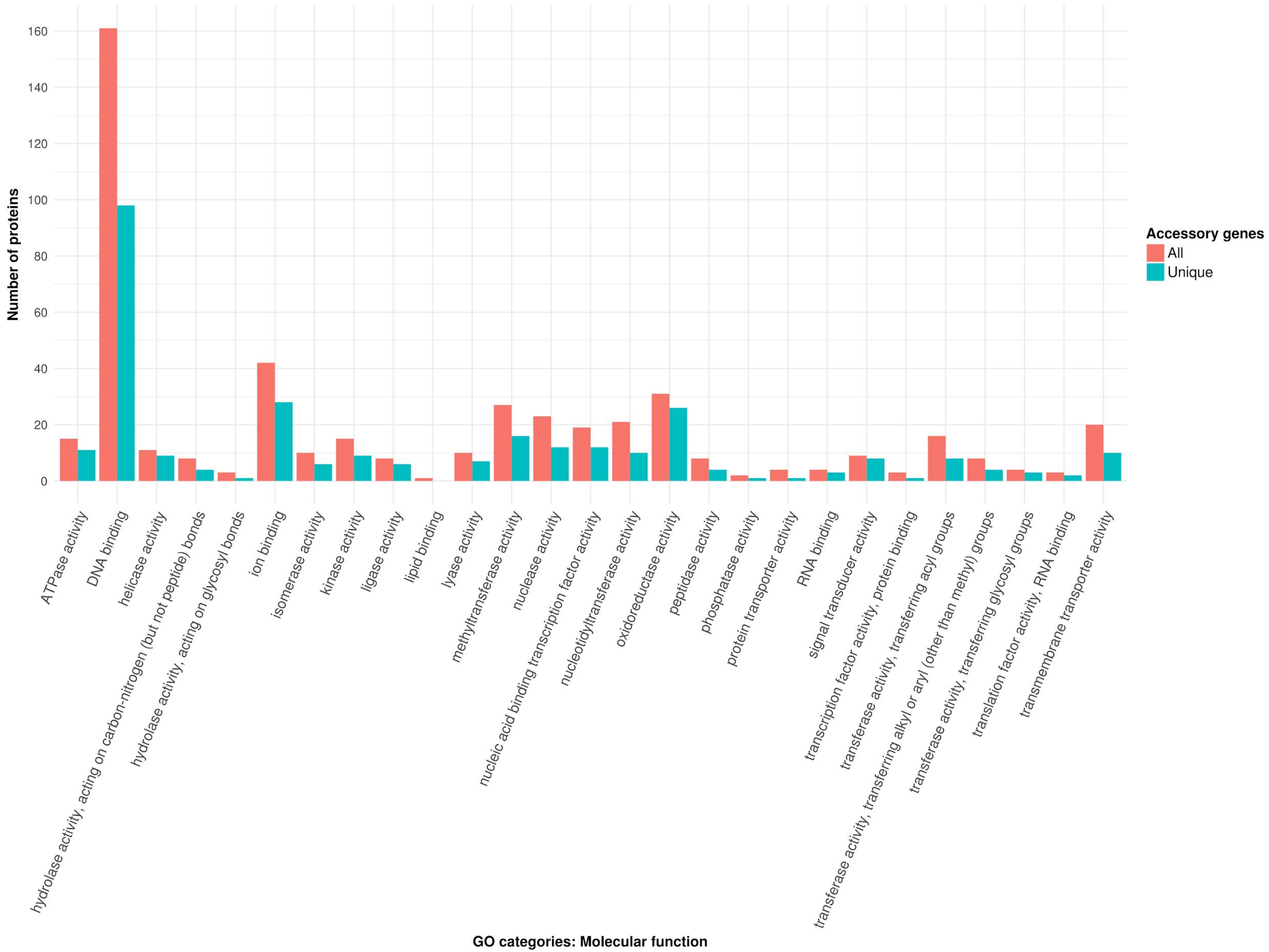

Figure S6. Functional annotation of the pBT2436-like megaplasmids family accessory genome

**B**

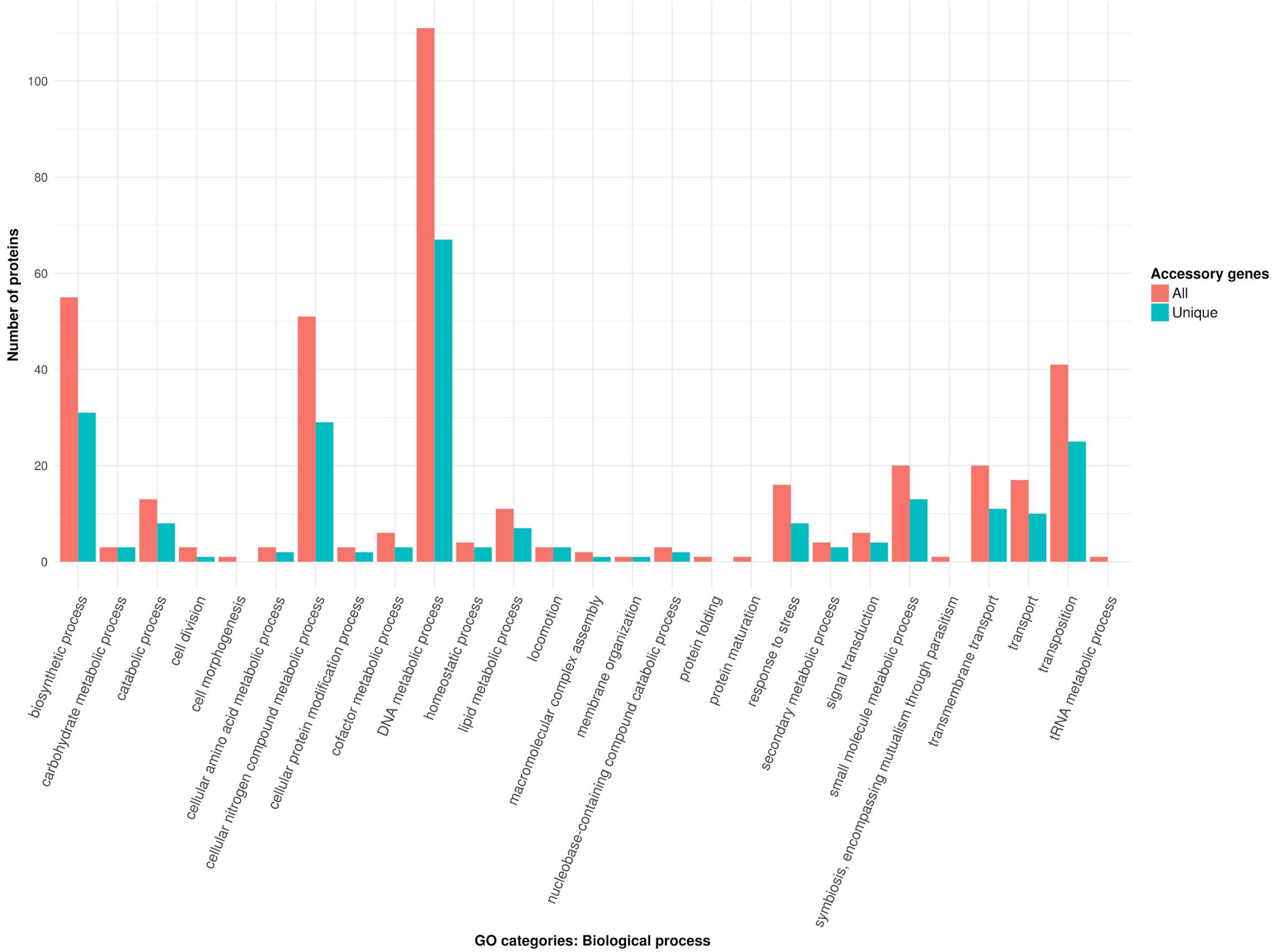

### Figure S6. Functional annotation of the pBT2436-like megaplasmids family accessory genome

**C**

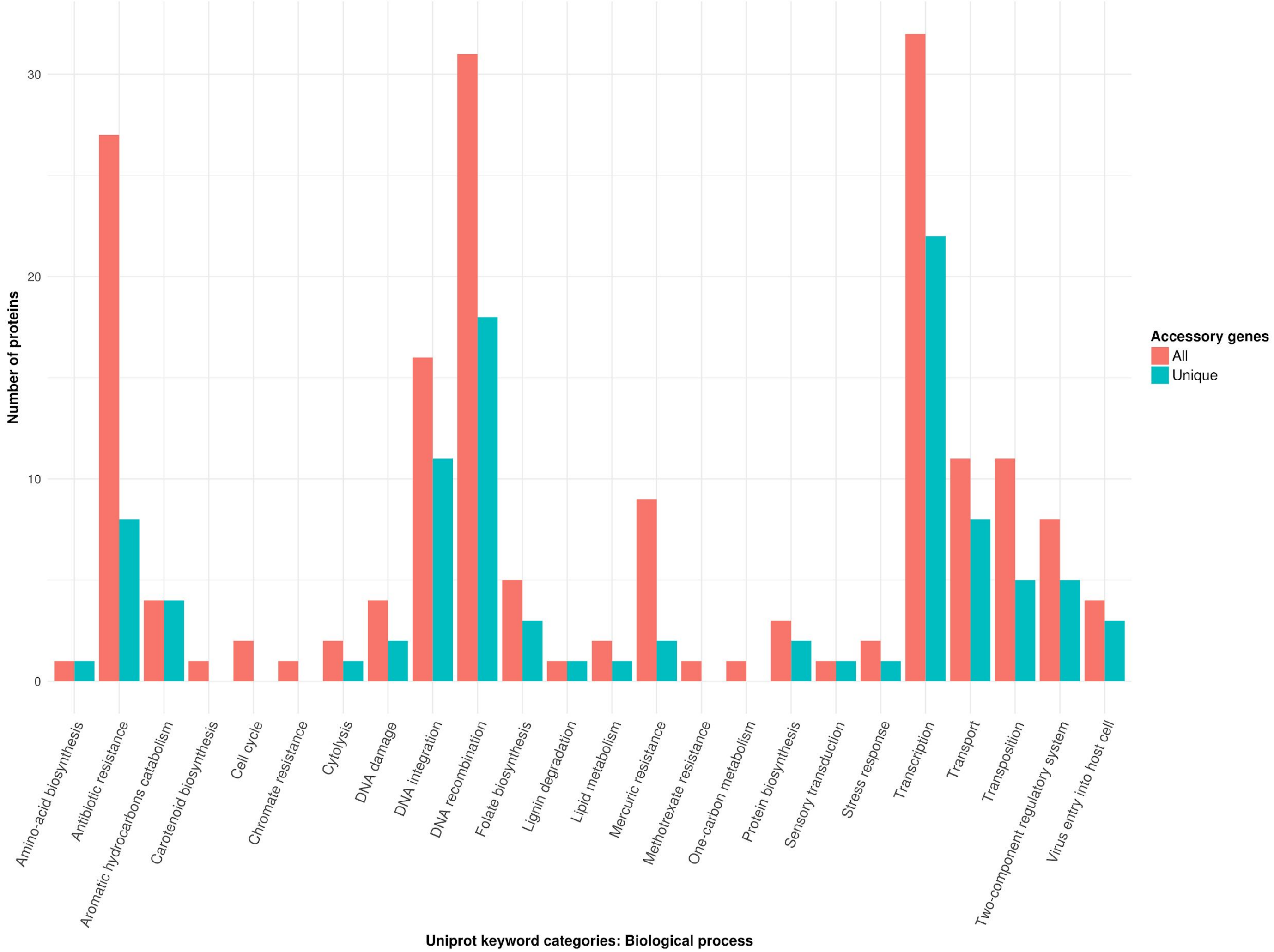

### Figure S7. Search of pBT2436-like megaplasmids in *Pseudomonas* genomes from the GenBank Assembly database

**A** Megaplasmids detection from contigs assembled from short-read sequencing data

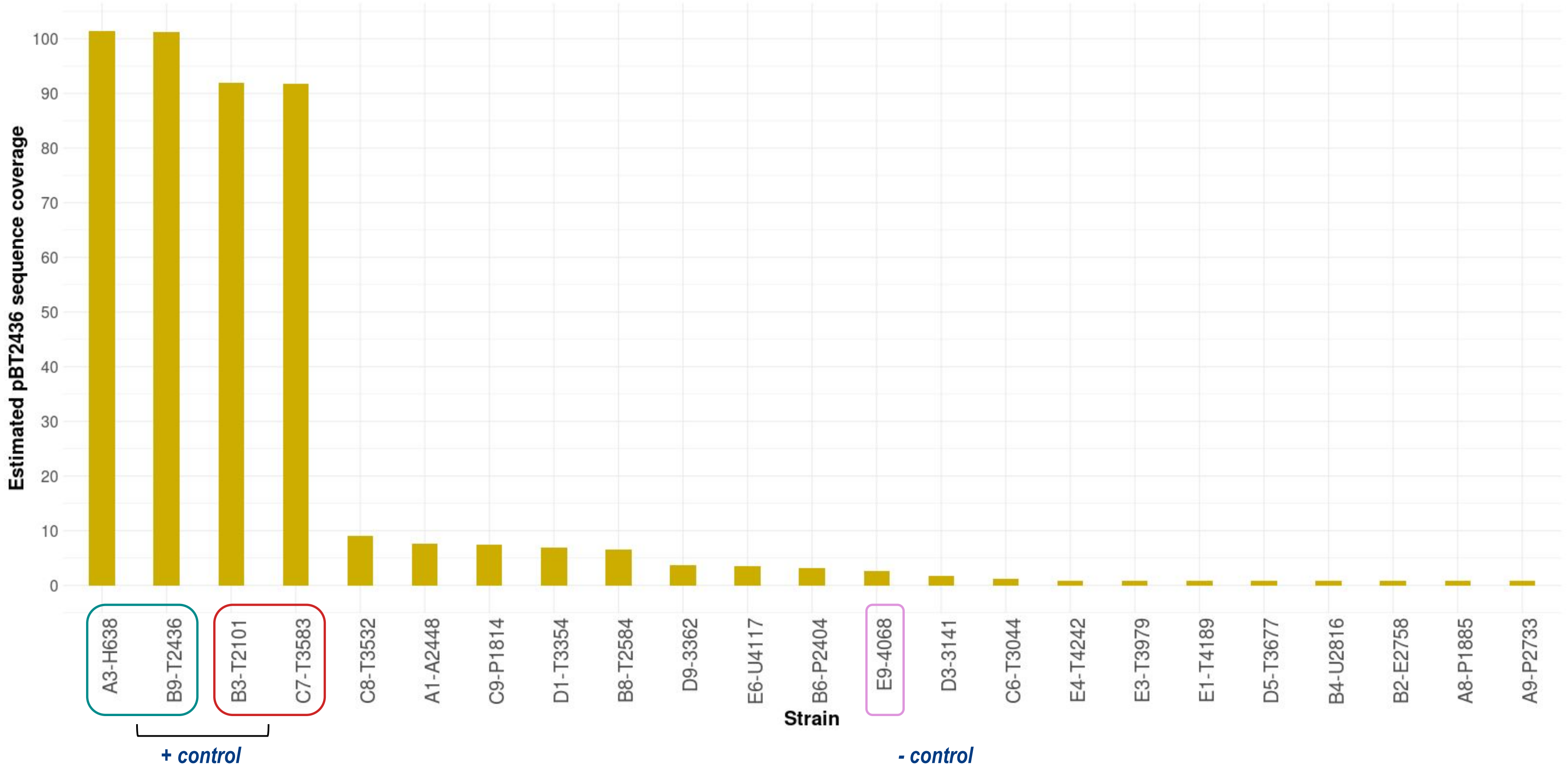

### Figure S7. Search of pBT2436-like megaplasmids in *Pseudomonas* genomes from the GenBank Assembly database

**B** pBT2436 coverage from alignments against contigs from *P. aeruginosa* Thai isolates genomes

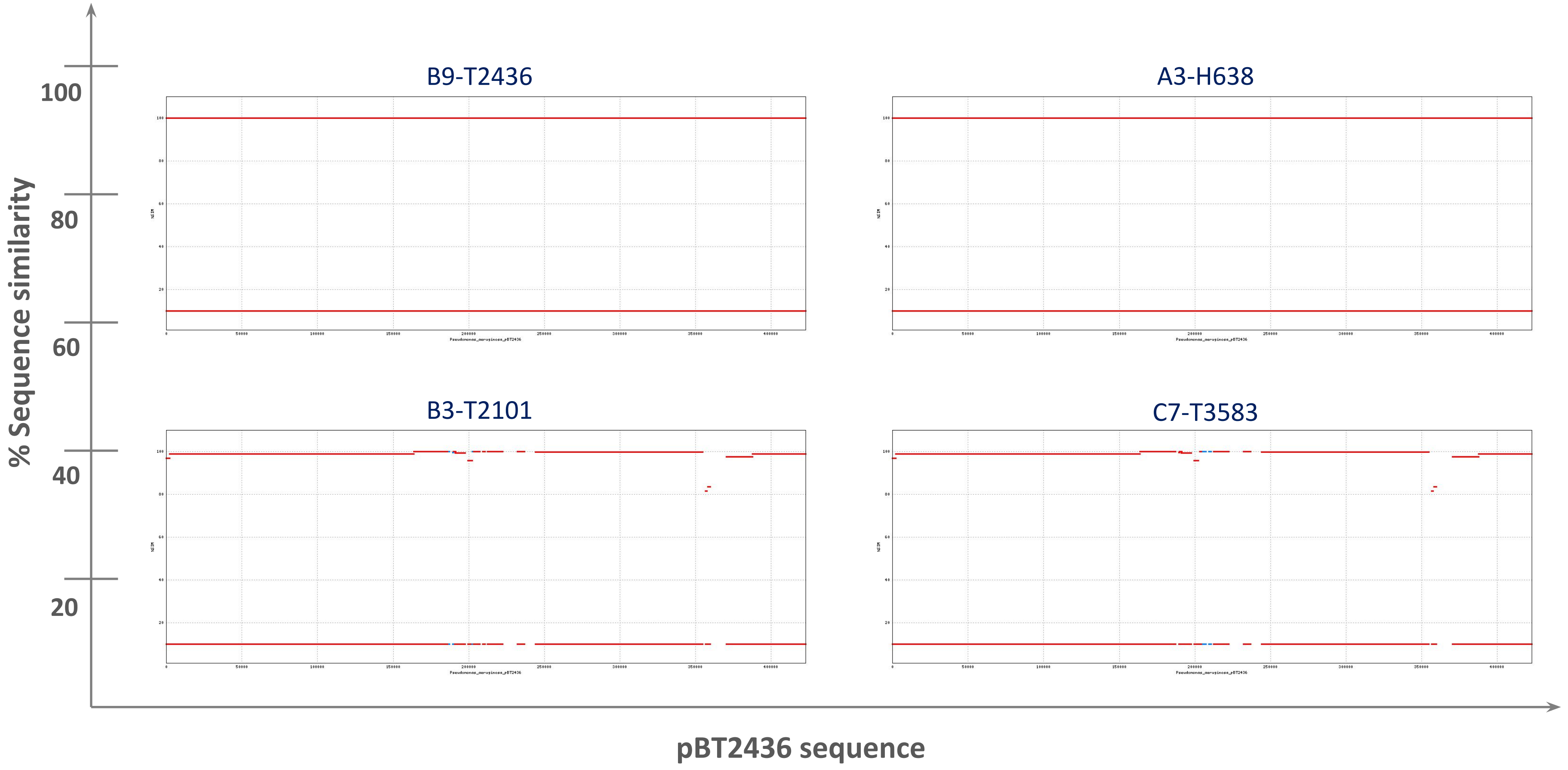

### Figure S7. Search of pBT2436-like megaplasmids in *Pseudomonas* genomes from the GenBank Assembly database

**C** Overlooked pBT2436-like megaplasmids in the *P. aeruginosa* genomes reported by Kos et al. 2015

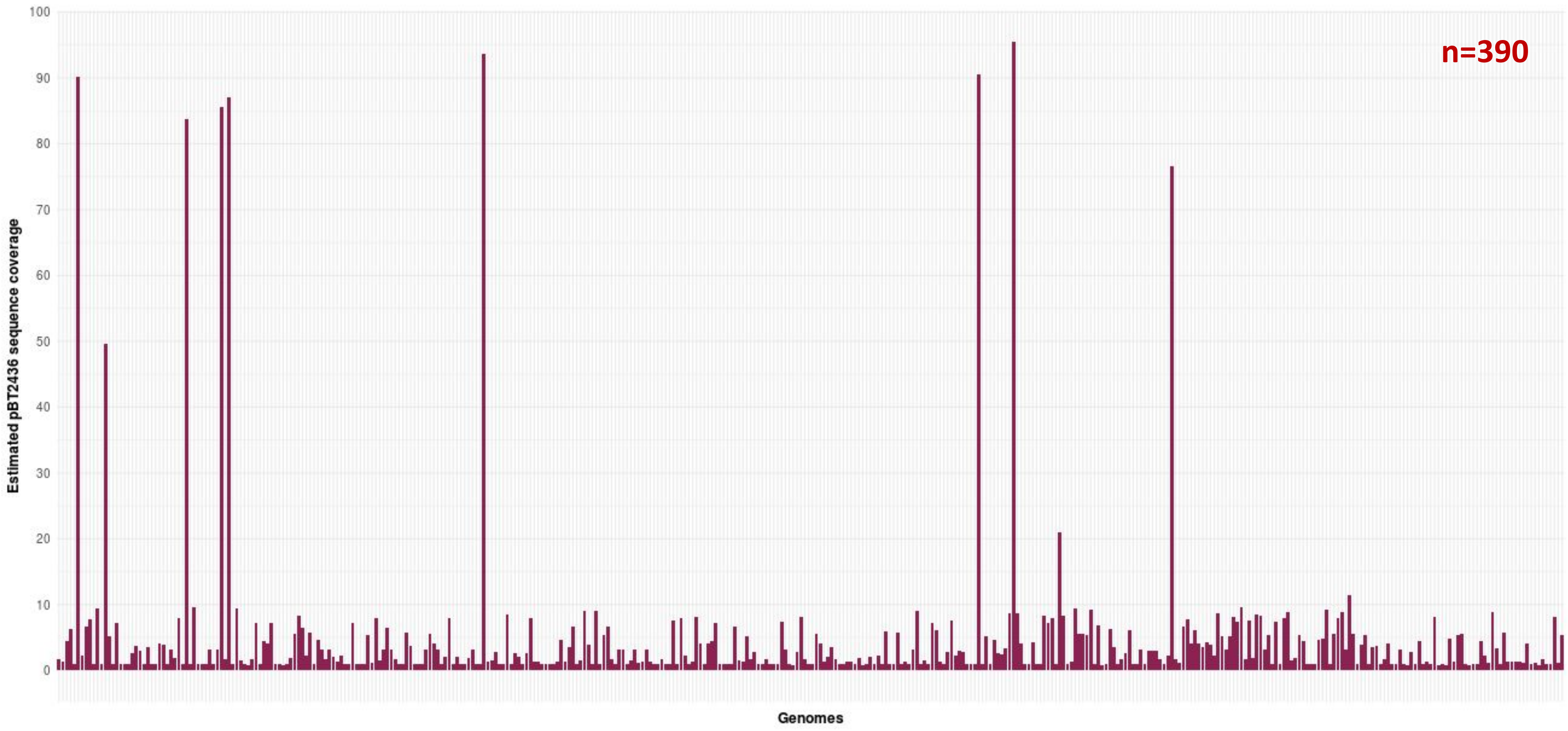

### Figure S7. Search of pBT2436-like megaplasmids in *Pseudomonas* genomes from the GenBank Assembly database

**D** pBT2436 coverage from alignments against contigs from *P. aeruginosa* genomes reported by Kos et al. 2015

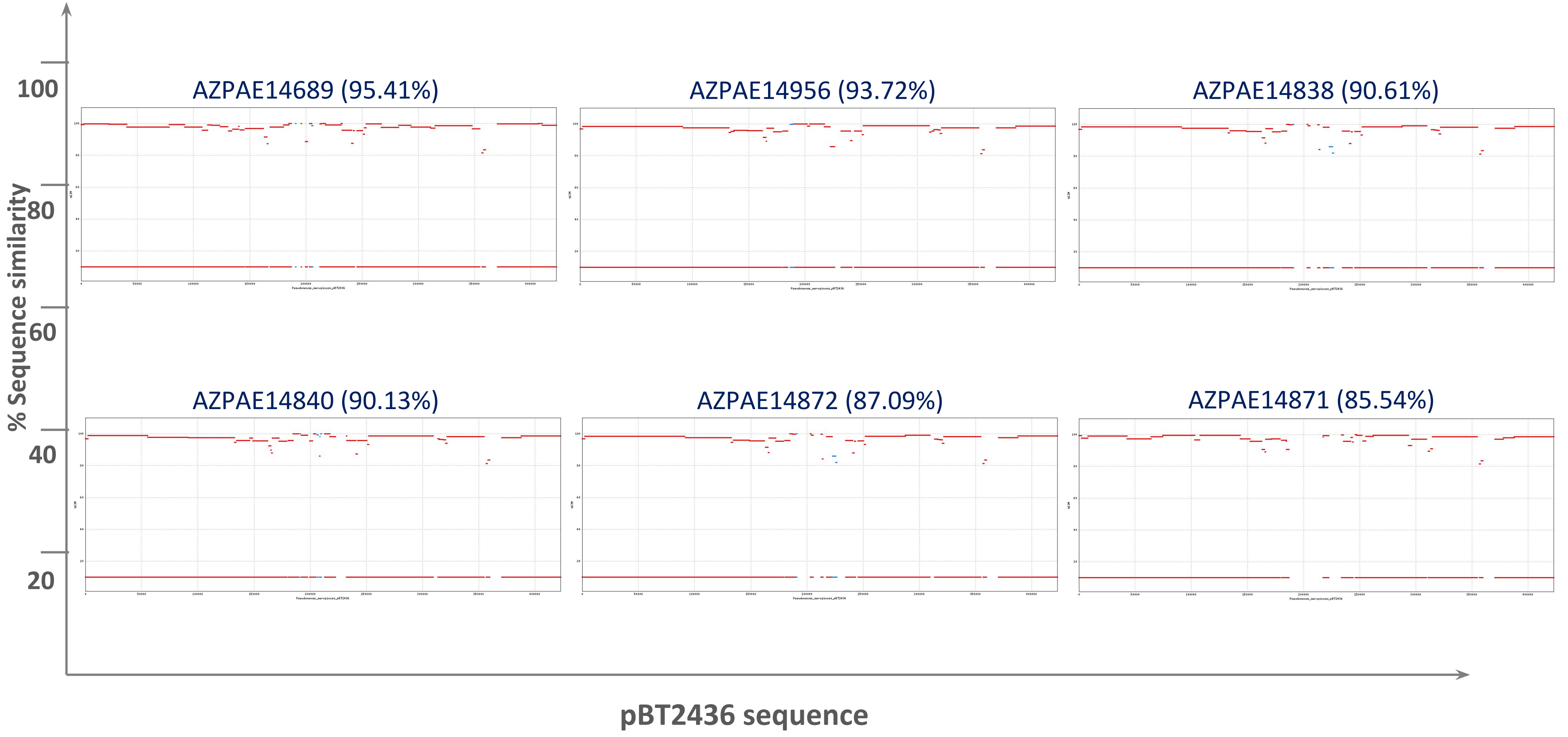

### Figure S7. Search of pBT2436-like megaplasmids in *Pseudomonas* genomes from the GenBank Assembly database

**D** pBT2436 coverage from alignments against contigs from *P. aeruginosa* genomes reported by Kos et al. 2015

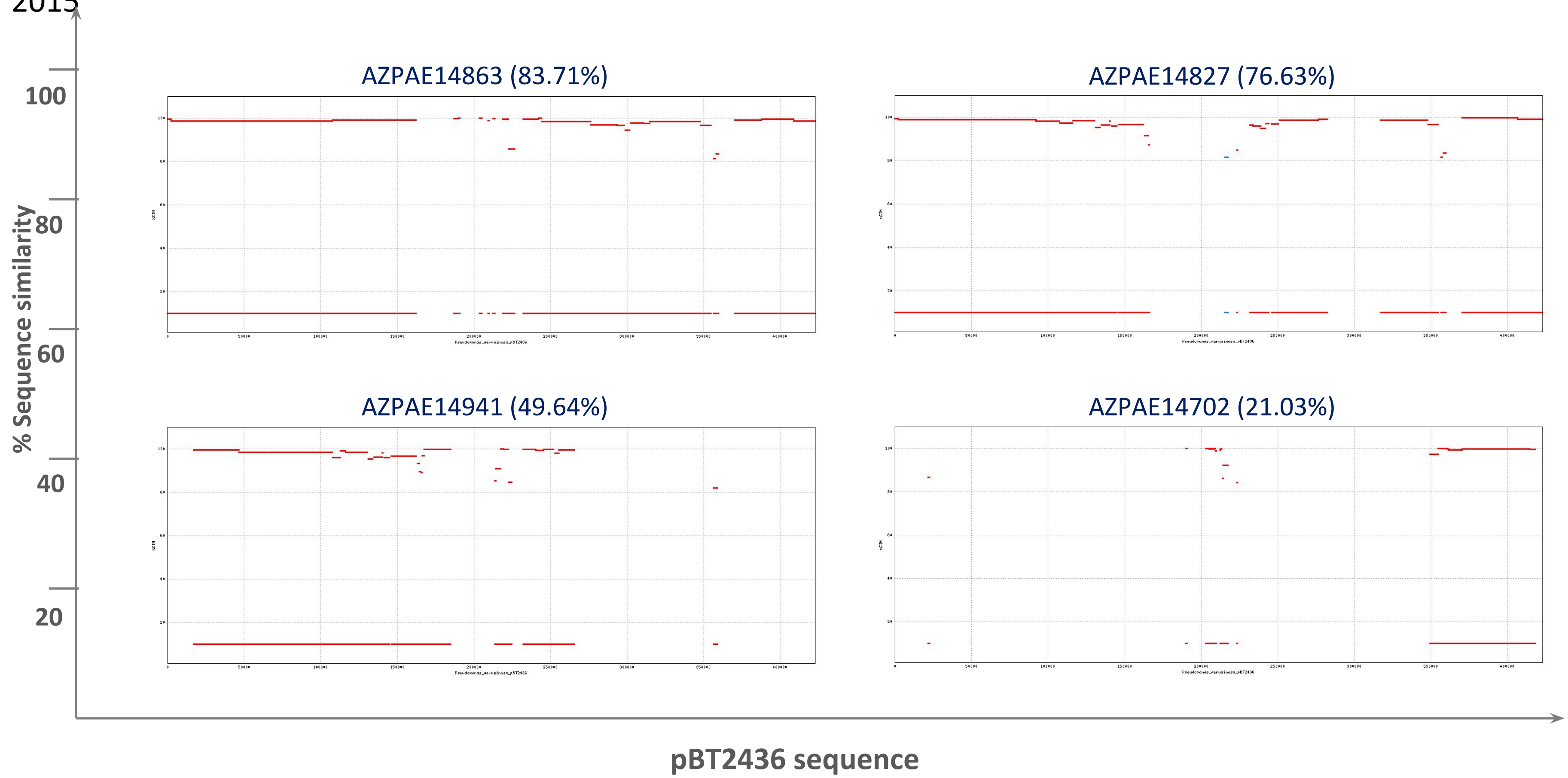
